# MPRAnalyze - A statistical framework for Massively Parallel Reporter Assays

**DOI:** 10.1101/527887

**Authors:** Tal Ashuach, David Sebastian Fischer, Anat Kreimer, Nadav Ahituv, Fabian Theis, Nir Yosef

**Affiliations:** Department of Electrical Engineering and Computer Sciences, University of California, Berkeley, California USA; Center for Computational Biology, University of California, Berkeley, California USA; Institute of Computational Biology, HelmholzZentrum Munchen, 85764 Neuherberg, Germany; TUM School of Life Sciences Weihenstephan, Technical University of Munich, 85354 Freising, Germany; Department of Bioengineering and Therapeutic Sciences, University of California San Francisco, San Francisco, California, USA; Institute for Human Genetics, University of California San Francisco, San Francisco, California, 94158, USA; Ragon Institute of MGH, MIT, and Harvard, Cambridge, MA, USA; Chan Zuckerberg BioHub, San Francisco, CA, USA

## Abstract

Massively parallel reporter assays (MPRAs) are a technique that enables testing thousands of regulatory DNA sequences and their variants in a single, quantitative experiment. Despite growing popularity, there is lack of statistical methods that account for the different sources of uncertainty inherent to these assays, thus effectively leveraging their promise. Development of such methods could help enhance our ability to identify regulatory sequences in the genome, understand their function under various setting, and ultimately gain a better understanding of how the regulatory code and its alteration lead to phenotypic consequence.

Here we present MPRAnalyze: a statistical framework dedicated to analyzing MPRA count data. MPRAnalyze addresses the major questions that are posed in the context of MPRA experiments: estimating the magnitude of the effect of a regulatory sequence in a single condition setting, and comparing differential activity of regulatory sequences across multiple conditions. The framework uses a nested construction of generalized linear models to account for uncertainty in both DNA and RNA observations, controls for various sources of unwanted variation, and incorporates negative controls for robust hypothesis testing, thereby providing clear quantitative answers in complex experimental settings.

We demonstrate the robustness, accuracy and applicability of MPR-Analyze on simulated data and published data sets and compare it against the existing analysis methodologies. MPRAnalyze is implemented as an R package and is publicly available through Bioconductor [1].

## Introduction

Enhancers are non-coding DNA sequences that contribute to the regulation of gene expression. Enhancers control the levels, timing and location of transcription, playing a crucial role in maintaining and determining cell identity and state. Sequence variants in enhancers can have significant consequences, as demonstrated by most disease-associated expression quantitative trait loci (eQTLs) identified through genome-wide association studies falling within non-coding regions of the genome [2].

Since enhancers regulate transcription by interacting with transcription factors and the transcriptional machinery, active enhancers tend to reside in areas of open chromatin. Additionally, enhancers have been shown to be marked by the histone modification H3K27ac [3, 4, 5, 6]. Assays that map these properties have been used extensively for genome-wide identification of active enhancers in various contexts [7, 8]. However, while these assays enable identification of candidate enhancers, they are limited to a binary view of enhancer activity, and are therefore insufficient to fully understand cis regulation of transcription. Functional assays are necessary to further our understanding of enhancers’ role in gene expression regulation. Reporter assays have been used to functionally annotate enhancers, by introducing fluorescent reporter constructs regulated by the enhancer of interest, but these assays have limited throughput and don’t scale to allow genome-wide functional annotations.

Recent advances in reporter assays address this issue, in a set of procedures denoted Massively Parallel Reporter Assays (MPRAs). These assays replace fluorescent reporters with sequence-based identifiers, denoted “barcodes”. Broadly, a synthetic construct that contains a minimal transcriptional structure is introduced into cells. Each such construct is generally composed of an enhancer of interest, a minimal promoter and a unique barcode. The synthetic enhancer is assumed to regulate the transcription of the barcode sequence similarly to how the native enhancer regulates the transcription of it’s target gene. The cells then undergo RNA and DNA sequencing to measure both RNA transcript counts and DNA construct counts for normalization purposes, and the RNA/DNA ratio is used to estimate the transcription rate. Relying on sequence-based identifiers allows using the vast combinatorial space of unique sequences instead of a limited set of fluorescent reporters, and leveraging next generation sequencing to measure the activity of thousands of enhancers in a given experiment.

MPRAs can be used to address several scientific questions. In mutagenesis experiments they are used to quantify the transcription rate of a variant, enabling a quantitative comparison to other variants, thereby measuring the effects of various mutations and alterations. Similarly, MPRAs can be used to classify enhancers as active: significantly affecting the native transcription rate of the promoter [9, 10]. In classification studies, control sequences are typically included to establish a baseline transcription rate for the minimal promoter used. Finally, MPRAs can be used for comparative analyses, comparing enhancer activity between different alleles [11, 12], tissues [13], or other conditions of interest. More complex experimental designs are also possible, for example measuring the interaction between alleles and condition [12], or measuring temporal behavior with time-series data [14].

Despite growing popularity of MPRAs, current studies have used various ad-hoc methods or methods that were not developed for MPRA data, such as DESeq2 [15], that relies on underlying assumptions that may not be true for MPRA data. Other MPRA analysis methods only address some of the types of questions MPRAs can address, such as QuASAR-MPRA [16] and mpralm [17] that only perform comparative analyses, and both rely on ratio-based summary statistics that limit the power of the analysis. In contrast, MPRAnalyze provides a general statistical framework that allows all uses of MPRAs to be address using a single model, leverages the unique structure and characteristics of MPRA data, and avoids relying on limited statistics or over-reaching assumptions.

## Results

MPRA data is produced from two parallel procedures: RNA-seq data from post-transduction cells measures the number of transcripts produced of each barcode, and DNA-seq data measures the number of construct copies of each barcode. Thus for each barcode in the experiment both DNA and RNA counts are observed, and the ratio RNA/DNA serves as a conceptual proxy for the transcription rate. However, both DNA and RNA measurements are products of sub-optimal and noisy procedures, an issue exacerbated by the unstable nature of a ratio: minor differences in the counts themselves can result in major shifts in the ratio, especially when dealing with small numbers. This problem can be handled by associating multiple barcodes with each enhancer, providing multiple replicates within a single experiment and a single sequencing library. This approach introduces an additional problem of how to properly summarize counts from multiple barcodes to a single transcription rate estimate of an enhancer, which is made difficult since transduction efficiency, while theoretically uniform across the different constructs, has a significant degree of variability in practice (Figure 1A). Two methods of summarizing are commonly used: the aggregated ratio, which is the ratio of the sum of RNA counts across barcodes divided by the sum of DNA counts across barcodes; and the mean ratio, which is the mean of the observed RNA/DNA ratios across barcodes. Both of these summary statistics have inherent limitations. The aggregated ratio loses the statistical power that multiple barcodes provide, and the mean ratio is highly sensitive to noise, as demonstrated by Myint et al. [17].

**Figure 1:**
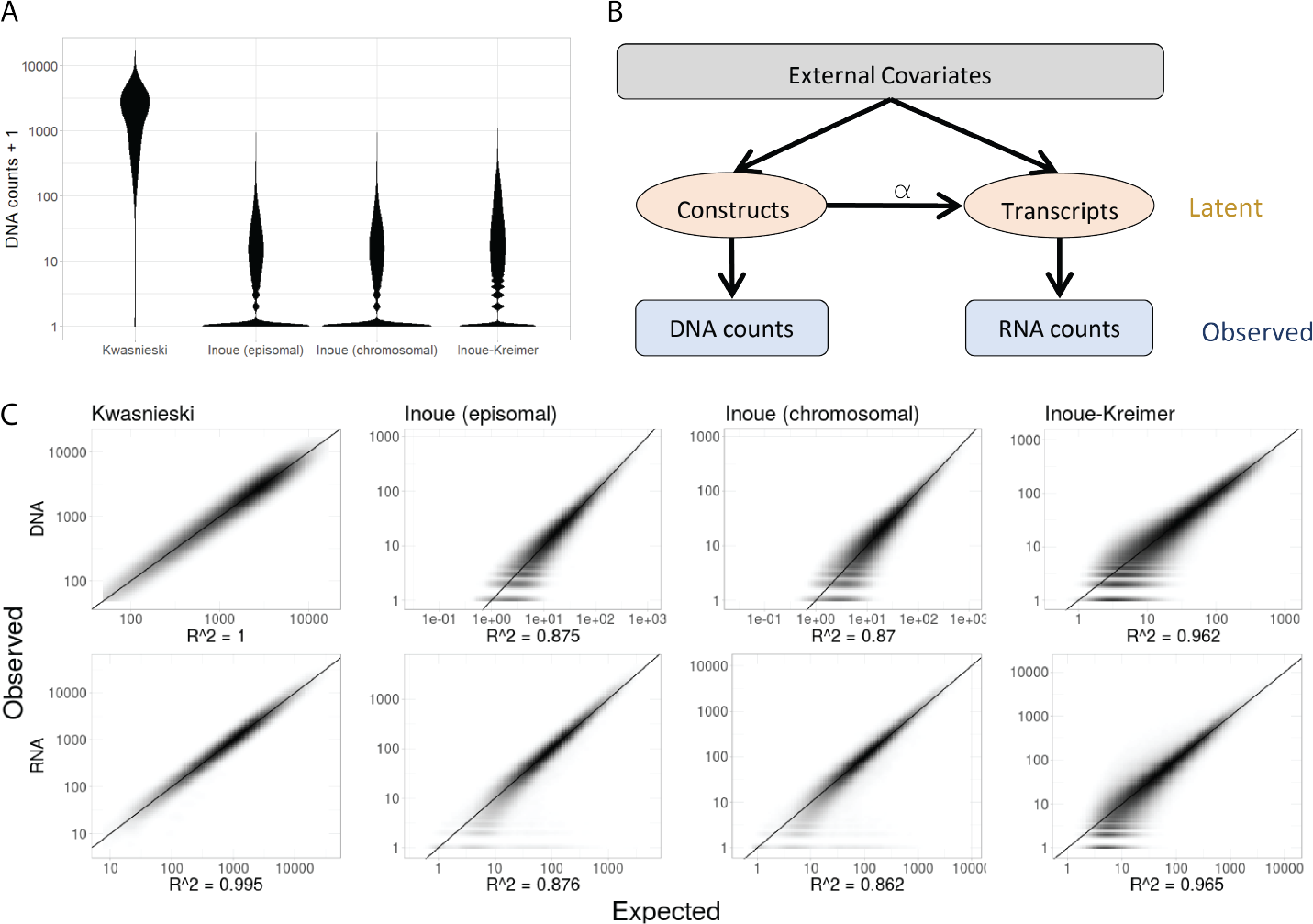
MPRAnalyze model properties and fit. **(A)** Distribution of construct abundances (DNA barcodes) across datasets, computed as the observed barcode count + 1 for visualization purposes. **(B)** A graphical representation of the MPRAnalyze model. External covariates (e.g conditions of interest, batch effects, barcode effects) are design-dependent; Latent construct and transcript counts are related by the transcription rate *α*. **(C)** Goodness of fit plots for both libraries across datasets. Expected counts were extracted from the fitted GLMs. MPRAnalyze’s model fits MPRA data well, with *R*^2^ > 0.86 across all datasets.

### 1 MPRAnalyze Model

We propose MPRAnalyze, a dedicated model for the analysis of MPRA data that uses a graphical model to relate the DNA and RNA counts, model the uncertainty in both libraries and take advantage of the unique structure and opportunities presented by MPRA data (Figure 1B). Our model relies on the assumption of a linear relationship between the RNA counts and the DNA counts: *RNA* = *DNA·α*, similarly to ratio-based approaches, with ‘*α*’ denoting the transcription rate. To account for the variability of barcode abundances as well as other covariates (conditions of interest, batch effects, etc), our model constructs two generalized linear models (GLM). The first GLM is the DNA model, which estimates the latent construct counts from the external covariates and the observed DNA counts. The second GLM, the RNA model, estimates the rate of transcription from the external covariates, estimated construct counts obtained from the DNA model, and observed RNA counts. Formally, for a given enhancer, we have two vectors observations: DNA counts 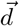 and RNA counts 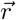. Then the MPRAnalyze models are:

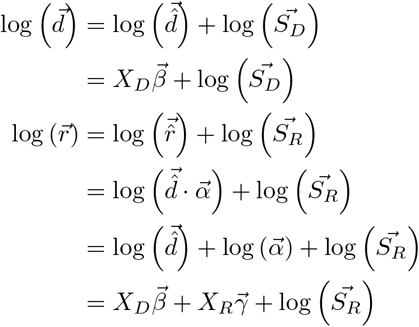

Where 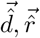 are the abundance estimates of the constructs and transcripts, respectively; *S*_*D*_, *S*_*R*_ are external normalization factors used to correct technical effects such as library size; 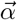 is the vector of transcription rate estimates, which can be either a single value if the analysis is looking for a single estimate (this would normally be the case for quantification and classification analysis), but can encode for multiple estimates for a single enhancer -for example if multiple biological conditions are analyzed simultaneously, the model can compute the *α* estimate for transcription rate for each condition; 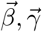 are the model parameters and *X*_*D*_, *X*_*R*_ are the design matrices, which encode the experimental design of the assay. Briefly, each column in the matrix corresponds to a single coefficient, and each row to a single sample. Numerical factors are incorporated as single columns, whereas for categorical factors (such as replicates or conditions of interest), each category has a separate coefficient and therefore a separate column, with one of the categories being absorbed as the reference (baseline) value, and the rest being treated as contrasts, and the values of the matrix being binary, determining the inclusion of each coefficient in the modelling of each sample (A simplified example is provided in Figure S1).

This formulation allows for straight-forward encoding of various covariates, and easily supports the common structure of MPRA experiments: multiple barcodes per enhancer, multiple replicates, and often multiple conditions analyzed simultaneously. This flexibility also allows for various covariates to only be modelled in one of the models, depending on the scientific question and experimental design. For instance, barcode-level effects should be incorporated into the DNA model to allow for proper normalization of the transcript counts, but should usually be excluded from the RNA model since we do not expect each barcode to result in a different transcription rate. Alternatively, in unpaired settings where the DNA sequencing was performed on pre-transduction libraries, there might not be separate DNA estimates for each condition being tested, in which case the conditions of interest would only be modelled in the RNA model, and excluded from the DNA model.

We optimize this model by maximizing the likelihood of the data using certain distributional assumptions. First, we assume that the latent construct counts, from which the observed DNA counts are sampled, are generated by a gamma distribution: 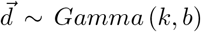. Second, we assume that the conditional distribution of the RNA counts follows a Poisson distribution: 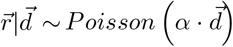, which results in a closed-form negative binomial likelihood for the RNA counts themselves: 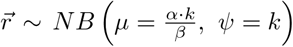. The negative binomial distribution is a common approximation of sequencing data, and indeed all datasets we examined have the negative-binomial characteristic quadratic relationship between the mean and the variance. This relationship is also observed for the DNA libraries, which is expected of Gamma-distributed data if the distribution’s shape parameter *k* ≈ 1 [Figure S2].

Our framework accounts for barcode specific effects and leverages them for increased statistical power while simultaneously benefiting from the robustness of aggregating information across barcodes. Since a standard for MPRA experimental design has yet to be formed, the nested GLM construction is flexible and can be easily extended to changing experimental designs. Our model is also highly interpretable, easily allowing for quantitative estimates of enhancer activity to be extracted, as well as differential activity to be tested directly using established statistical tests. Our framework also explicitly leverages negative controls when available, either to establish the null distribution in classification analyses or to correct for systemic bias in comparative analyses [see Methods].

To characterize the properties and evaluate the performance of the MPR-Analyze model, we compared MPRAnalyze’s performance and the properties of our model to other previously used and newly developed methods, using both simulated data and a set of four MPRA datasets detailed in Table 1. These datasets were chosen for representing a diversity of MPRA procedures (episomal or lentiviral integration, DNA sequencing from pre- or post-transduction), study focus (quantification, classification and comparative analyses), and experimental design (number of barcodes per enhancer). Note that only a subset of features of the Kwasnieski datasets are used (Only the weak and strong categories were used), and only a subset of samples in the Inoue-Kreimer dataset are used (only timepoint T0 in the quantification and classification analyses, and only T0 and T72 in the comparative analyses).

**Table 1:** MPRA datasets used for evaluation of MPRAnalyze throughout the paper.

| Dataset | Description | Integration | #Enhancers | #Controls | #Barcodes |
| --- | --- | --- | --- | --- | --- |
| Kwasnieski | Testing ENCODE putative weak/strong enhancers in K562 cells. [10] | Episomal | 1200 | 568 | 4 |
| Inoue (episomal) | Candidate liver enhancers in HepG2 cells without genomic integration. [9] | Episomal | 2338 | 102 | 100 |
| Inoue (chromosomal) | Candidate liver enhancers in HepG2 cells with lentivirus-based genomic integration [9] | Lentiviral | 2338 | 102 | 100 |
| Inoue-Kreimer | time-series MPRA during neural differentiation induction in human ESCs. [14]. For convenience, referred to as *Inoue-Kreimer*. | Lentiviral | 2464 | 200 | 90 |

When fitted to MPRA data, we found that the MPRAnalyze model is able to properly capture the characteristic of the data and provides a good fit across all datasets (*R*^2^ > 0.86 for all datasets, figure 1C). To examine the validity of assuming the DNA counts are gamma-distributed, random data was generated from the fitted DNA GLM and the residuals were compared with the residuals of the observed counts. Quantile-based comparisons shows that the residuals are generated by similar distributions [Figure S3], indicating that this assumption does not significantly distort that distribution of the observed data.

### 2 Quantification

We set out to examine the properties of MPRAnalyze’s estimate of transcription rate, denoted ‘alpha’, and compare it to the naïve ratio-based summary statistics. Overall the three estimates are largely in agreement (Pearson’s *r* > 0.9 across datasets, Figure S4), demonstrating that alpha is indeed capturing the correct signal.

To examine the accuracy of the estimates, we used the negative control sequences included in some of the datasets. These are assumed to have an identical transcription rate induced by the minimal promoter with no additional enhancer activity. We examined the variance of the estimates on these sets. In the Kwasnieski dataset, which has a limited number of barcodes, the three estimates all had low variance. In the barcode-rich datasets, alpha was clearly more consistent across the negative controls than the other two estimates, with the aggregated ratio being the least consistent (Figure 2A).

**Figure 2:**
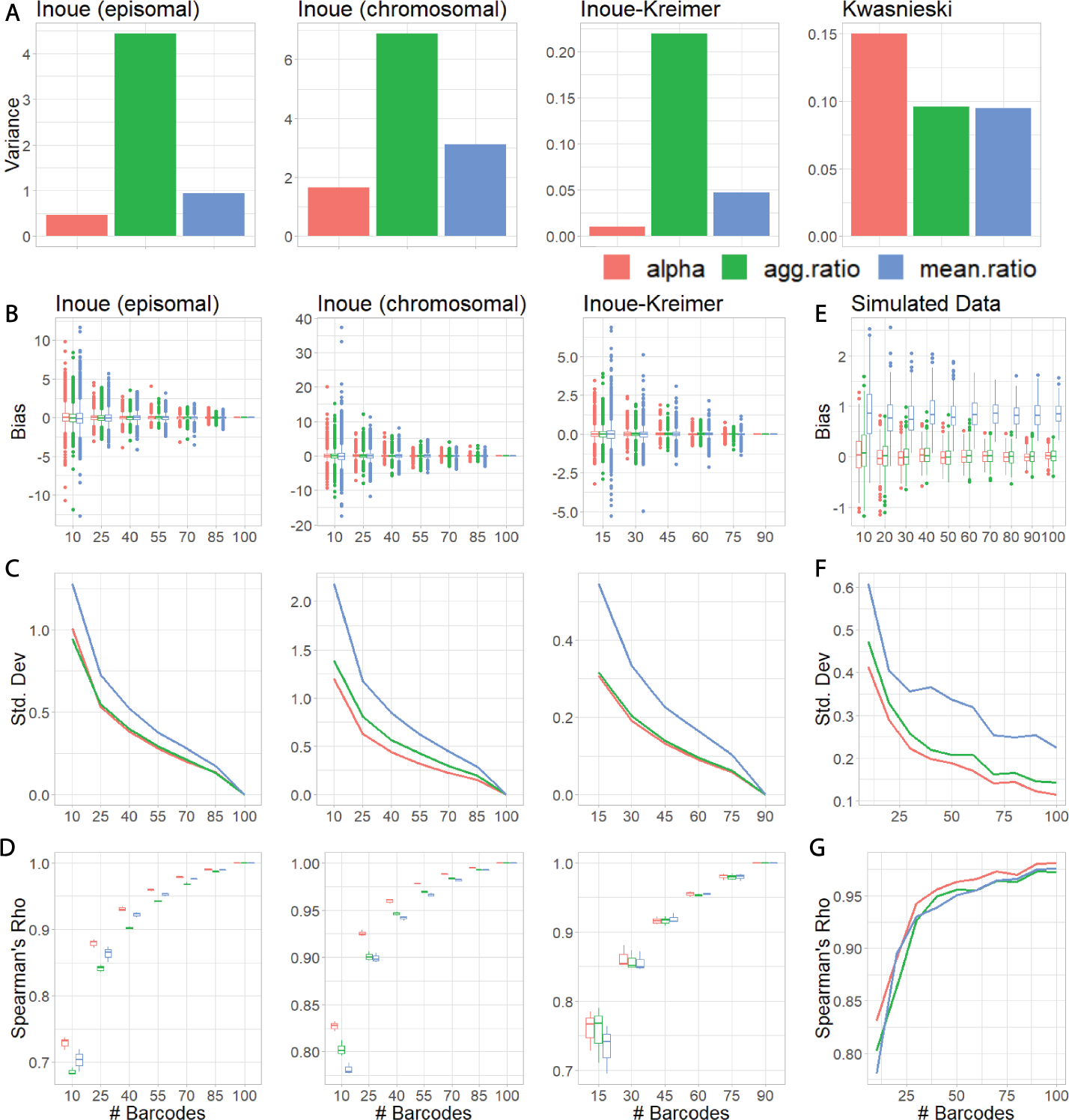
Comparison of MPRAnalyze’s *α* estimate of transcription rate with the naive ratio-based estimates. **(A)** The variance measured among estimates of negative-control enhancers in each dataset (these are assumed to have an identical transcription rate). **(B-D)** Barcodes were subsampled and quantification was recomputed based on the partial data to measure the effect of barcode number on estimate performance [See methods for further subsampling details]. Analyses were performed using the full-data estimate as the ground truth. **(E-G)** MPRA data was simulated to provide an actual ground truth. In each case we measured the bias (*estimate − truth*)(**B,E**); the standard deviation 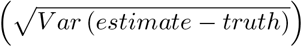 (**C,F**); and the Spearman correlation between the estimates and the ground truth **(D,G)**.

We then set out to explore the effect of the number of available barcodes on the performance of the estimates. Using the barcode-rich datasets, barcodes were subsampled at various rates and estimates were recomputed for each enhancer (3 independent samples per enhancer per barcode rate). Using the full-data estimates as a baseline, we found that subsampling barcodes does not result in a systemic bias in any of the estimates (Figure 2B). Expectedly, all estimates showed reduced variance with increased barcodes, with the mean ratio underperforming the other two estimates, and alpha having a similar or lower variance than the aggregated ratio (Figure 2C).

In many cases the goal of quantifying enhancer activity is to rank and compare different enhancers, as in mutagenesis experiments. To compare the stability of the ordering of enhancers, the Pearson correlation was computed between the estimates in each sub-sample to the estimates of the full data. Alpha has either similar or higher correlation than both naive estimates across datasets and barcode abundance (Figure 2D).

Noting that these analyses are limited by a lack of ground truth, MPRA data was then simulated by generating random coefficients and using the same nested GLM construction as described above to generate samples. To avoid biasing the results, samples were generated with a log-normal noise model, instead of the default Gamma-Poisson convolutional model MPRAnalyze uses [methods]. We generated 101 enhancers with gradually increasing transcription rates (from 2 to 3, in 0.01 steps). The analyses above were repeated with the simulated data. We found that while the measured bias was indeed not influenced by the number of barcodes, the mean ratio displayed a significant amount of bias compared with both alpha and the aggregated ratio (FIgure 2E). Similar to the real data results, we found alpha has lower variance than both naive estimates, and higher correlation with the true transcription rates (Figure 2F-G).

Overall, alpha is as or more stable and accurate as the aggregated ratio when barcode information is limited, and is more consistent across similarly-behaving enhancers than both the aggregated and the mean ratio.

### 3 Classification

MPRA-based classification of active enhancers has previously been done by comparing the ratio-based estimates of candidate enhancers to the control set [9, 10], an approach that suffers from the limitations of the summary statistics demonstrated above. Other studies performed this analysis using DESeq2 [15], a differential expression analysis (DEA) method, by treating the DNA and RNA libraries as two conditions and looking for “differentially expressed” enhancers [11]. However, the method relies on an implicit assumption that the majority of features do not display differential behavior, a valid assumption for DEA that does not hold for classification of MPRA data, in which the candidate enhancers are often explicitly selected as sequences that are likely to be active.

MPRAnalyze performs classification of active enhancers by testing each enhancers alpha estimate against a null distribution describing the null transcription rate induced by the minimal promoter without enhancer activity. When negative controls are available, they are used to estimate the null distribution. When they are not available, MPRAnalyze relies on a conservative assumption that the mode of the distribution of transcription rate estimates is the center of the null distribution, and that values lower than the mode all belong to the null. Thus, MPRAnalyze estimates the null by locating the mode and using only the values lower than the mode to estimate the variance. Each enhancer’s alpha values are then tested by computing Median-Absolute-Deviation (MAD) scores, median-based variants of Z-scores that are less sensitive to outliers.

To assess MPRAnalyze’s performance in classification analyses we compared 6 methods: MPRAnalyze with and without controls; empirical p-values computed using the naive ratio estimates; and DESeq2 in either full mode (barcode-level data) or collapsed mode (summing across barcodes within each batch). DESeq2 hypothesis testing was performed using an asymmetric alternative hypothesis, only looking at enhancers that were more active in the RNA library than in the DNA library.

We examined the fraction of enhancers that were significantly active (FDR < 0.05) in each dataset, stratified by group: negative controls, candidate enhancers and positive controls when available. Expectedly, empirical p-values from the naive ratio estimates show a clear lack of power, only being comparably sensitive in the Inoue-Kreimer dataset. Both DESeq2-collapsed and MPRAnalyze without controls have inflated rates of activity in the Kwasnieski datasets, however overall both modes of DESeq2 and both modes of MPRAnalyze have reasonable results across datasets. Overall, MPRAnalyze and DESeq2 seem to find comparable numbers of active enhancers, with no clear advantage to either method [Figure 3A].

**Figure 3:**
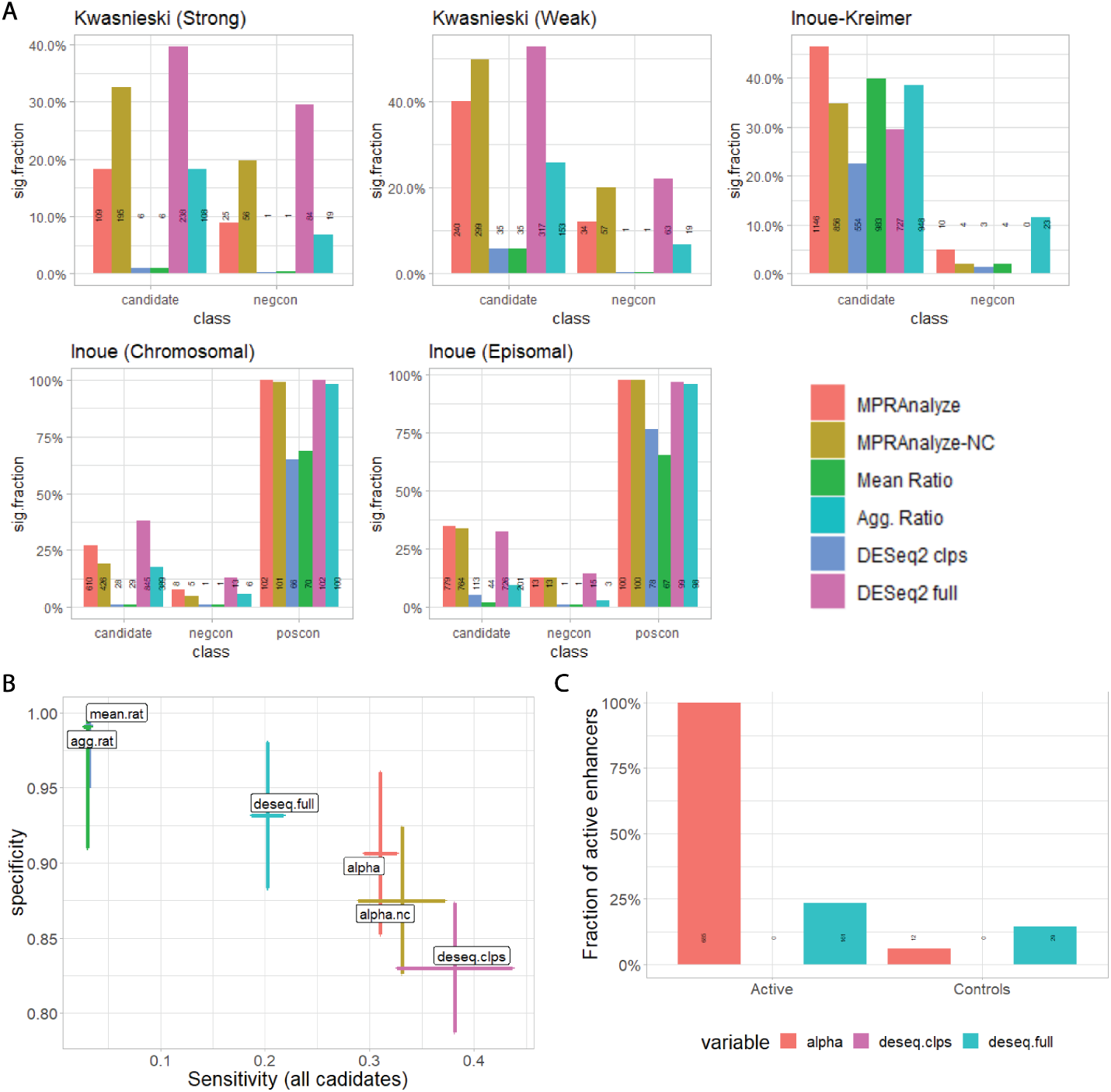
Classification analysis comparisons. (**A**) fraction of enhancers identified as significantly active (BH-corrected *P* < 0.05) by method and class of enhancer. MPRAnalyze results both in control-based (red) and no-controls (orange) modes; empirical p-values based on the mean ratio (blue) or aggregated ratio (green); DESeq2 results in collapsed mode (barcodes are summed within each batch, purple) or full mode (full data, light blue). Absolute number of active enhancers is displayed on the bars. (**B**) Precision-Recall curve. Precision is based on performance on the negative controls, Recall is based on the total population of enhancers, assuming all candidates are active. Error bars are ± the standard deviation of these measures across datasets. (**C**) Fraction of active enhancers detected after re-running the analyses on 685 enhancers from the Inoue-Kreimer dataset that were identified as active by MPRAnalyze (regular mode) and both DESeq2 modes, and the 200 controls from the same dataset. MPRAnalyze recapitulates the same results, finding that 100% of the candidates are active, whereas DESeq2 full only identifies 161 (23.5%) and DESeq2 collapsed completely fails to identify any active enhancers.

However, only looking at the fraction of enhancers that pass a threshold misses the overall statistical behavior of the method. We therefore examined the full p-value distribution of each method within each dataset, finding that despite comparable rates of enhancers found statistically significant, the MPRAnalyze model is far better calibrated to MPRA data. When negative controls are used, MPRAnalyze-generated p-values follow the theoretical behavior of the statistic: uniform on null-generated data, low values otherwise. When negative controls are not used, MPRAnalyze has some deviations from behavior, emphasizing the importance of including negative controls in classification experiments, to properly characterize the null behavior. Both MPRAnalyze modes have significantly higher statistical power, demonstrated in the distribution of p-values on the positive controls included in the Inoue datasets. Conversely, DESeq2 p-values do not follow the theoretical behavior of a well-calibrated test, instead mostly having a concentration of high and low p-values, both among candidates and controls [Figure S5].

We hypothesized that these calibration issues are partly explained by the asymmetric alternative hypothesis test we used when running DESeq2. When directly comparing p-values from MPRAnalyze to the competing methods, we indeed found that overall the scores are correlated, and that the abundance of high-valued p-values are mostly among enhancers that MPRAnalyze and the ratio-based p-values do not reject the null for, indicating that DESeq2 views these as “down regulated”, meaning enhancers who’s expression in the RNA library is lower then DESeq2 model expects it to be based on the DNA library counts [Figure S6].

One of the major aspects of the DESeq2 model is the dispersion shrinkage mechanism that the model uses. This is common practice among differential expression methods, and includes pooling information across all features included in the dataset (genes for RNA-seq, candidate enhancers in MPRA). Since RNA-seq is a genome-wide assay, the set of measured features across different experiments can be assumed to remain stable, if not necessarily constant. This does not hold true for MPRA experiments, in which the number and composition of the measured regulatory sequences is curated according to the specific goals and context of the given experiment. We therefore hypothesized that DESeq2-based classification would be highly dependent upon the composition of enhancers included in the analysis. To demonstrate this, we re-ran the analysis on the Inoue-Kreimer dataset, but only included the 200 control enhancers and 685 enhancers that were classified as active by MPRAnalyze, DESeq2-full and DESeq2-collapsed. This simulates a scenario in which an identical experiment was performed, producing the same data, but included fewer enhancers that were selected with higher degree of certainty of their activity. Since MPR-Analyze only pools information across enhancers when correcting for library size, we expected it to recapitulate the original results and indeed MPRAnalyze finds all candidates are significantly active. However, DESeq2-full only identifies 161 (23.5%) of the candidates as active, and DESeq2-collapsed finds no active enhancers at all [Figure 3C]. These results are not surprising, as the high abundance of activity would shift DESeq2’s estimate of the null behavior, whereas MPRAnalyze avoids using the entire population to estimate the null. This reveals an inherent reproducibility issue in using differential expression analysis designed for RNAseq to perform MPRA classification.

### 4 Comparative Analysis

Another common use for MPRAs is comparative studies, looking for differential behavior of enhancer induced transcription between conditions. These are often comparisons between different tissues, stimuli or comparing different alleles of an enhancer sequence [11, 13]. More complex experimental settings are also possible, e.g measuring temporal activity as in the Inoue-Kreimer data [14], or the interaction between differential allele activity and GATA1 presence in Ulirsch et al [12].

MPRanalyze’s model construction is based on generalized linear models, and as such is highly flexible and extendable to various experimental designs. Performing differential activity analysis in the MPRAnalyze framework can be done in two straight-forward ways: first, since MPRAnlyze optimizes the model using likelihood maximization, any single hypothesis that can be encoded in a generalized linear model can be tested using the likelihood ratio test. This includes complex hypotheses that are not encoded in a single coefficient. Additionally, in simple two-condition designs, or in cases where multiple contrasts are compared to a single reference (e.g multiple different stimuli compared against the unstimulated behavior), the model coefficients can be extracted from the RNA model and tested using a Wald test. Both options are supported in the current implementation of MPRAnalyze, however results in this paper are based on the likelihood ratio test option.

When performing differential activity analysis in MPRA data, it can be important to account for possible systemic bias, for example if the cell types being compared have different inherent transcription rates. In RNA-seq experiments this issue is usually resolved via library size correction, but with MPRA this is not necessarily sufficient, and may actually introduce further bias. This is because for the library size to properly correspond to inherent bias in the data, either the vast majority of features must be non-differential, or the differential signal must be symmetric. Neither of these assumptions necessarily hold for MPRA data, since the features are curated and vary between different experiments. An MPRA can be designed with all features being up-activated (more active in the contrast condition than the reference), and in which the vast majority of features are indeed differentially active. To address this issue, MPRAnalyze utilizes negative controls in the data to define the null differential behavior. This is done be fitting a separate, joint model for the controls, in which each control enhancer has a distinct DNA model but they all share a single RNA model, essentially finding the common activity pattern across the conditions. The model coefficients of this joint model are then incorporated into the model fitted for each candidate enhancer as additional correction factors [see Methods].

Alternative methods have been developed to address this question. QuASAR-MPRA [16] was designed specifically for allelic-comparisons and uses a beta-binomial model and mpralm [17] which is a general differential-activity tool designed for MPRA which fits a linear model. Both methods use summary statistics and do not include barcode-level information in their model. Mpralm enables using either the aggregated ratio or the mean ratio as the statistic, and therefore inherently suffers from the limitations of these statistics described above. QuASAR-MPRA, similar to MPRAnalyze, models the DNA and RNA separately, but it does so on the sum of counts across all barcodes in each condition, collapsing the data into a single measurement.

To compare the performance of MPRAnalyze to the above methods, we used the Inoue-Kreimer dataset and extended the subset of samples we used to include both T0 and T72 timepoints (0 and 72 hours into neural induction of hiPSCs). We then compared enhancer activity between the different timepoints, using the four methods: MPRAnalyze, mpralm in both aggregated ratio and mean ratio modes, and QuASAR-MPRA.

The distribution of p-values [Figure 4A] shows that overall MPRAnalyze and both modes of mpralm are well calibrated, following the expected mixture of uniform and low values among candidates, and showing slight inflation but overall uniform behavior among the controls. Conversely, QuASAR-MPRA seems poorly calibrated on both candidates and controls, recapitulating the results described by Myint et al [cite]. Indeed, QuASAR-MPRA only identified 2 candidates as significantly differential (BH-corrected p-values < 0.05).

**Figure 4:**
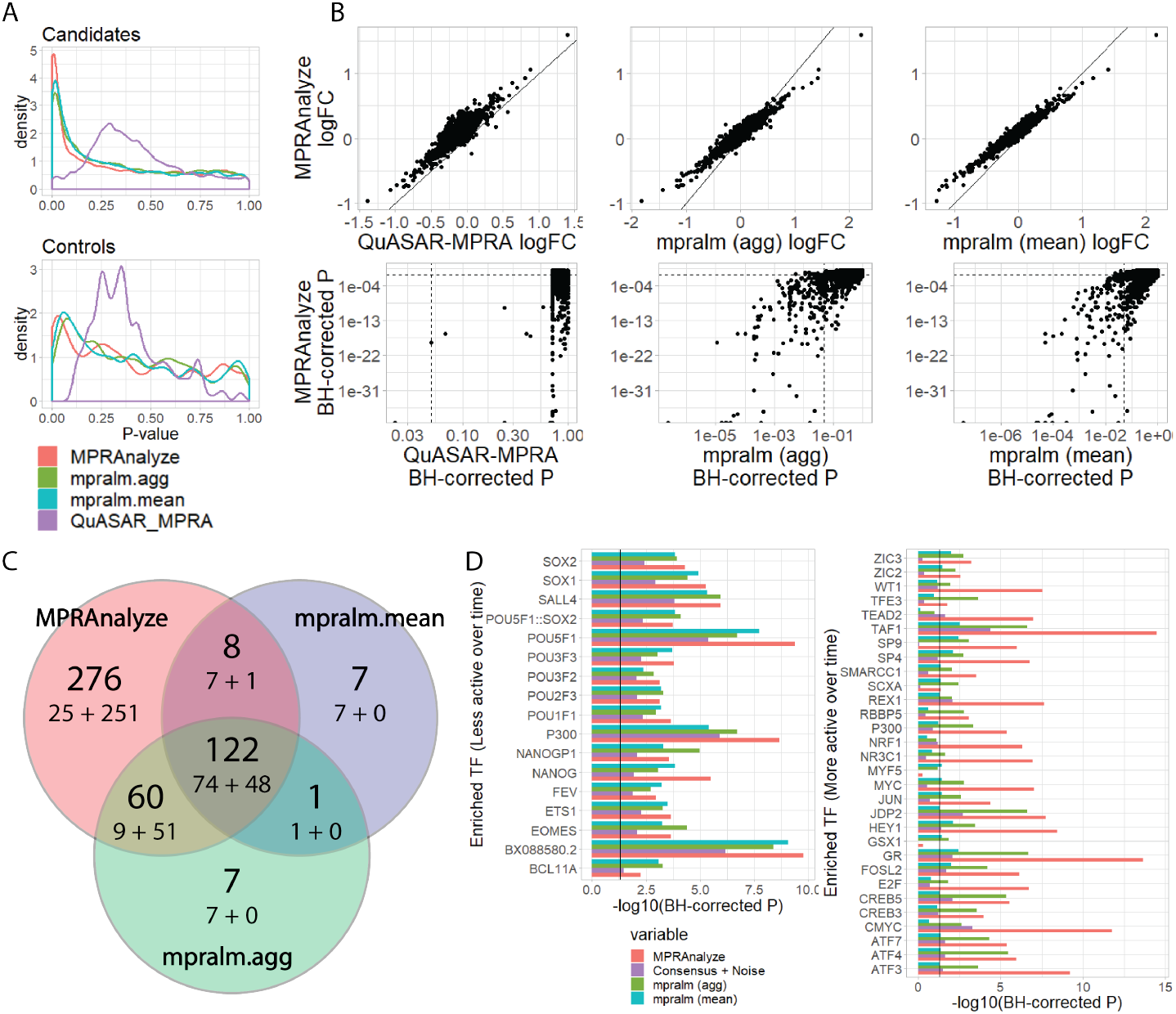
Comparative analysis results of comparing timepoint 0h to 72h in the Inoue-Kreimer dataset. (**A**) P-value distributions of candidates (top) and controls (bottom). QuASAR-MPRA is poorly calibrated, whereas MPRAnalyze and both mpralm modes follow the theoretical behavior (mixture of uniform and low values). (**B**) Direct comparison of MPRAnalyze to competing methods. Top panels show the biological effect size (log Fold-change); Bottom panels show the statistical significance (BH-corrected P; dotted lines are 0.05 threshold). (**C**) Venn diagram for MPRAnalyze and mpralm (both modes). The numbers in each area are (top) the total number of enhancers in the area, and (bottom) the number of decreasing-activity enhancers (left) + and increasing-activity enhancers (right). (**D**) Enrichment of transcription factor binding sites in differentially active enhancers as determined by each method. Solid line represents threshold of 0.05. (see Methods for further details).

We then examined the agreement among the methods by comparing the log Fold-Change estimates produced by each method. For QuASAR-MPRA we used the *betas.beta.binom* value, which is the logit transformation of the allelic skew, a proxy for biological effect size. Overall all four methods are well correlated (Pearson’s r ¿ 0.84 across all pairs), so similar biological effect is being observed by all methods. MPRAnalyze is more conservative than both mpralm modes, and has overall a positive skew compared with QuASAR-MPRA [Figure 4B].

We then compared MPRAnalyze’s statistical effects to the other methods by looking at BH-corrected p-values [Figure 4B]. Both modes of mpralm are correlated with MPRAnalyze among statistically significant candidates, but demonstrate reduced statistical power. QuASAR-MPRA is expectedly uncorrelated, since the model is not statistically well calibrated.

Further examination of the results excluded QuASAR-MPRA since it did not identify enough differentially activated candidates for the subsequent analyses. After filtering the results to only include candidate enhancers that are also classified as active in at least one of the conditions (BH-corrected *p* < 0.05, using MPRAnalyze’s classification method, since mpralm does not support classification analysis), we found that MPRAnalyze identifies a higher number of significantly differentially active candidates than mpralm in either mode [Figure 4C]. Interestingly, mpralm in aggregate mode finds a roughly balanced number of enhancers that are increasing (99) and decreasing (91) in activity, and in mean mode finds more decreasing (89) than increasing (49), while MPRAnalyze finds far more increasing (351) than decreasing (115) enhancers. However, the experimental design of the selected enhancers in the Inoue-Kreimer dataset explicitly favored sequences that corresponded to increase in gene expression over the course of differentiation, therefore MPRAnalyze’s up-skewed results are expected, and better correspond to the dataset design.

We then explored the set of candidates that were detected by each method. For each candidate sequence potential transcription factor binding sites (TFBS) were identified [see Methods]. The set of differentially active enhancers we divided to decreasing and increasing, then within each set each transcription factor was tested for enrichment (hypergeometric test, BH-corrected *p* < 0.05). To narrow down the results, we examined the union of top 15 most enriched transcription factors by each method [Figures 4D, full results at Supp.]. Additionally, we examined a *consensus*+*noise* option, wherein we took the consensus set of enhancers for which all methods agree and added artificial counts to match MPRAnalyze number of significant enhancers. These counts were taken from the remaining population and are proportional to the fraction of enhancers that contain a binding site to some factor, to simulate enrichment inflation that doesn’t reflect true biological signal [See methods]. Among decreasing-activity enhancers, on which the three methods are largely in agreement, we get an expected similarity of enrichment scores across transcription factors, with MPRAnalyze outperforming the other methods in two of the core pluripotency factors (NANOG, POU5F1). Among increasing-activity enhancers, in which the differences between the three methods are more profound, we find a higher variability of enrichment scores across the top-enriched factors. Overall, mpralm in *mean* mode fails to capture many of the enriched transcription factors captured by the two other methods, only finding 23 enriched transcription factors (compared with 106 and 195 found by mpralm *aggregate* and MPRAnalyze, respectively) and having diminished statistical power. When comparing MPR-Analyze to mpralm *aggregate*, we find overall agreement on the set of enriched factors, with MPRAnalyze consistently having increased statistical power with few exceptions. This increase in power cannot be explained simply by MPRAnalyze’s detection of more enhancers, as can be seen by the *background* + *noise* results, demonstrating that MPRAnalyze’s results reflect true biological signal. Notable disagreements between the methods include TEAD2 and NRF1 that are enriched in MPRanalyze’s results but not in either mode of mpralm. Both factors have been implicated in neurogenesis by previous studies [18, 19], and show increased expression in the later time frame [Figure S7]. In the other direction, MYF5 and GSX1 are enriched according to mpralm but not enriched according to MPRAnalyze. However, when examining mRNA levels of these factors, both factors have low expression levels, below their characteristic expression levels in tissues they are known to be active in (based on GeneCard[20] reported levels), making them less attractive candidates of driving differential gene expression.

Overall, MPRAnalyze identifies a similar biological signal to the competing methods, with increased statistical power which allows for more nuanced results.

## Discussion

Massively Parallel Reporter Assays are a powerful technique for functional characterization of enhancer activity in a high-throughput manner. MPRAs can be used to directly quantify enhancer-induced activity and identify active enhancers, compare regulatory activity of different alleles, elucidate regulatory grammar via mutagenesis studies, and compare enhancer activity between conditions. Complex experimental design can include interaction studies, studying how sequence changes affect differential enhancer activity [12], or identifying temporal activity patterns in time-course MPRA data [14].

Since MPRAs are still a nascent technology, they often vary in experimental design. While MPRAnalyze can analyze any MPRA dataset, the method benefits from certain experimental decisions that are generally recommended but not always leveraged in other analyses. First, pairing the DNA and RNA libraries by extracting DNA from the same post-transduction libraries that the RNA libraries are extracted from, avoids introducing further experimental noise into the data, and enables MPRAnalyze to better fit and relate the two models to increase accuracy of estimating nuisance factors. Additionally, as demonstrated in our results, increasing the number of available barcodes can greatly reduce the measured noise and increase performance of all methods. Finally, the inclusion of negative control sequences allows explicit modeling of the null behavior and avoids relying on assumptions that may bias the results and prevent proper interpretation of them. The curated nature of MPRA datasets makes negative control a valuable and often crucial aspect of properly interpreting the results.

MPRAnalyze is the first method to emerge that offers a robust statistical framework that enables analyzing data for all major uses of MPRAs in a unified model. MPRAnalyze models noise in both DNA and RNA libraries and uses a powerful nested GLM design to control barcode-specific effects and leverage the multiplicity of barcodes for increased statistical power. The method is highly flexible and allows various complex study designs to be tested in a straight-forward manner. Additionally, MPRAnalyze avoids relying on population-level properties in the analysis, instead leveraging negative controls when available to establish null behaviors.

## Supporting information

Supplemental Methods - model

---

**Supplemental Figure 1:**
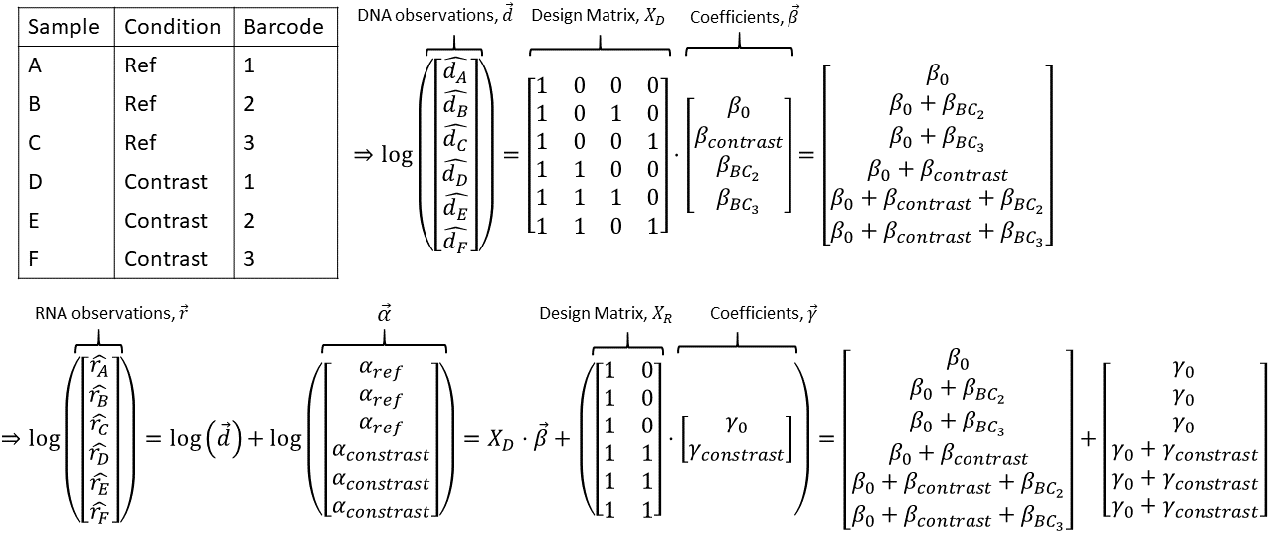
A simplified example of the MPRAnalyze model: two conditions are tested with three barcodes in a paired experiment (each DNA observation has a corresponding RNA observation). No replicates or external normalization factors are included in this design to maintain simplicity. The DNA’s model estimation of the latent DNA count, computed as 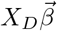 is included in the RNA model. The *α* estimates of transcription rate can be extracted from the model as: *α*_ref_ = *e*^*γ*_0_^, *α*_ref_ = *e*^*γ*_0_+*γ*_contrast_^. Note that while modeling the barcodes in the RNA model is possible, the result will be a separate *α* estimator for each barcode, which is usually not desired. Barcode-level information is therefore only incorporated into the RNA model via the nested DNA model.

**Supplemental Figure 2:**
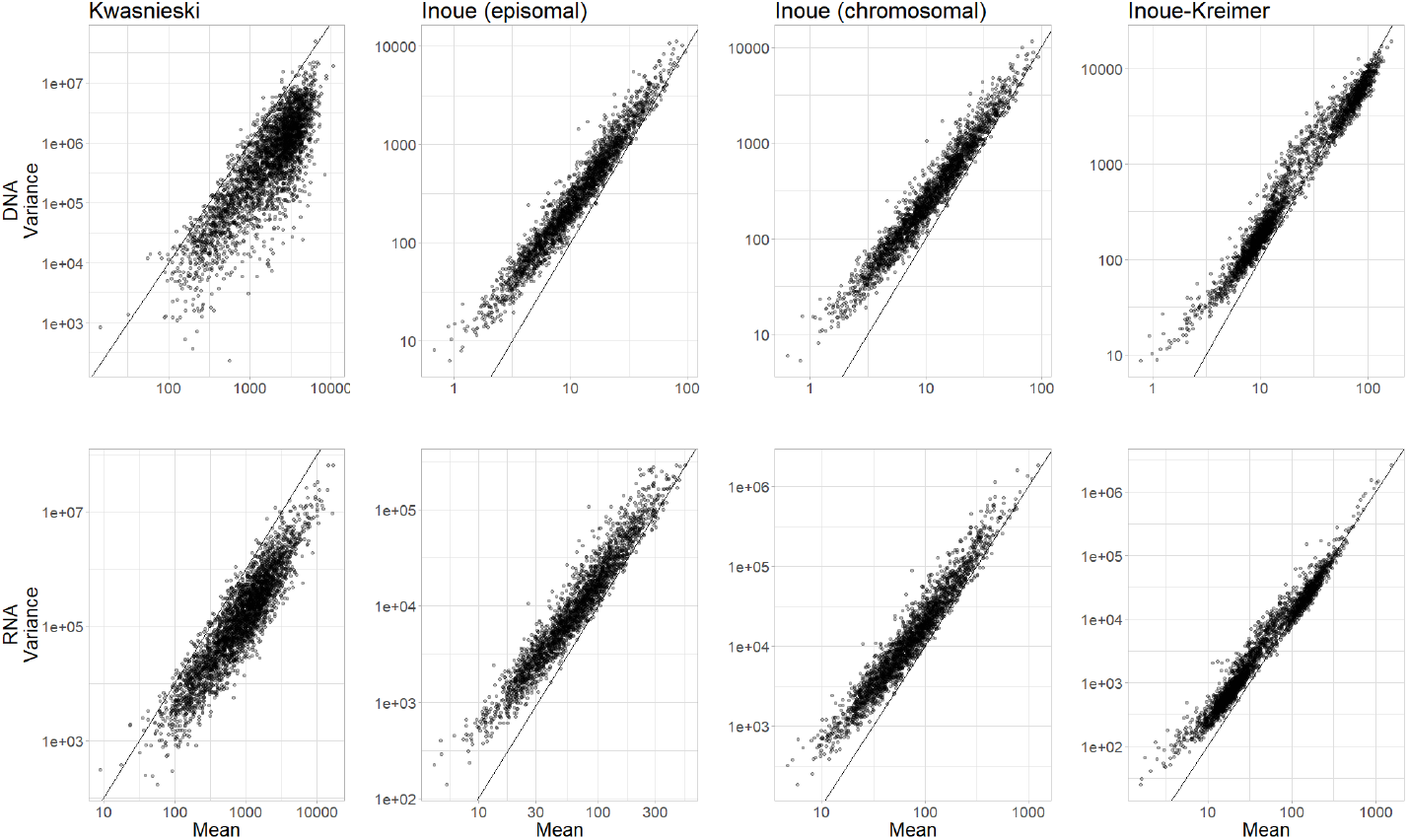
Relationship between the mean and variance of the counts measured for each enhancer. Reference line (slope = 2) is a quadratic relationship.

**Supplemental Figure 3:**
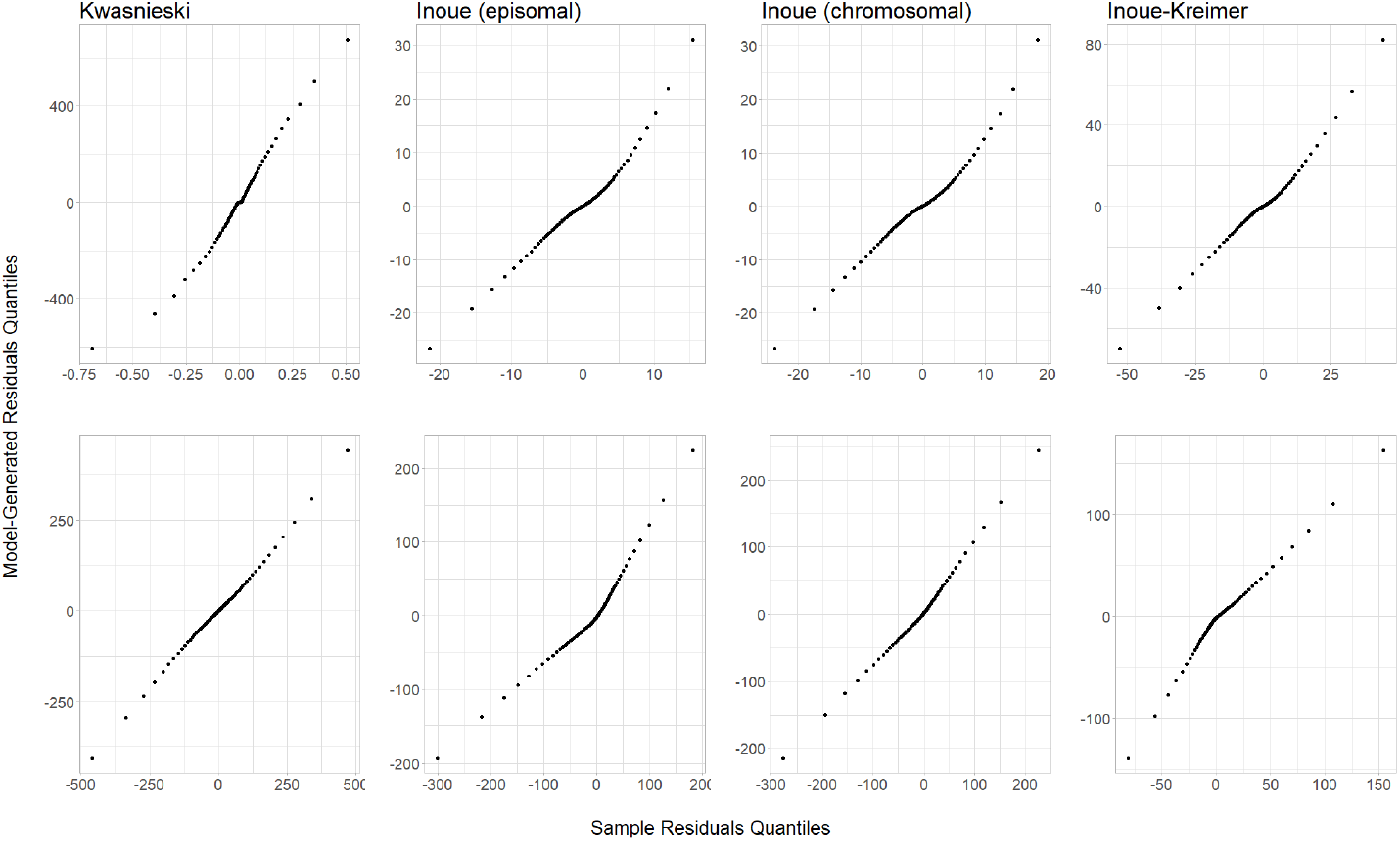
Comparison between model residuals from the observed counts of each dataset, and residuals from data generated by Gamma (for DNA) and Negative Binomial (for RNA) using the model parameters. Quantile-quantile comparisons indicate that the observed noise and the generated noise follow similar distributions.

**Supplemental Figure 4:**
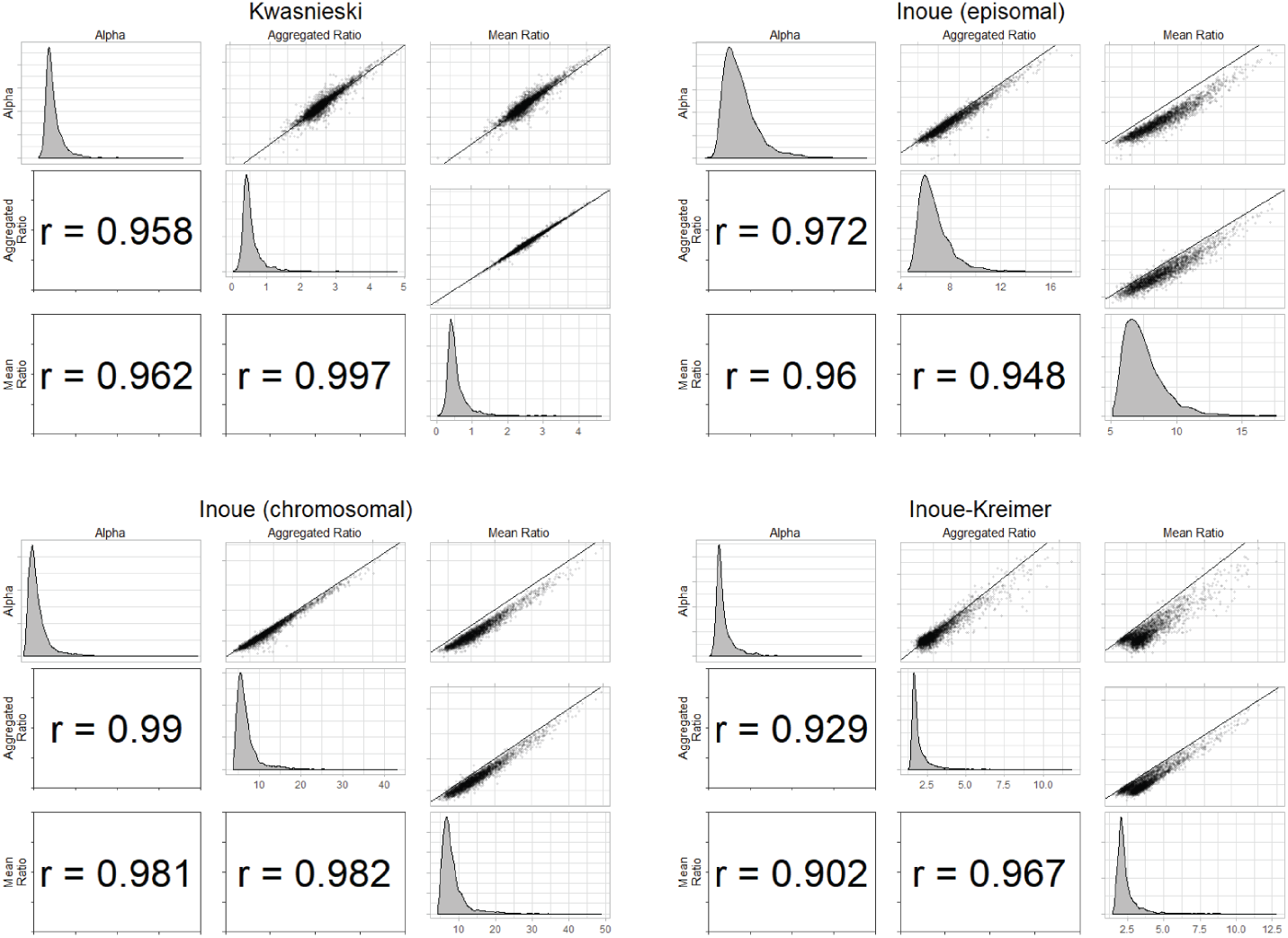
Correlations of MPRAnalyze’s *alpha* estimate with the naive ratio-based estimates. Correlations are Pearson’s r.

**Supplemental Figure 5:**
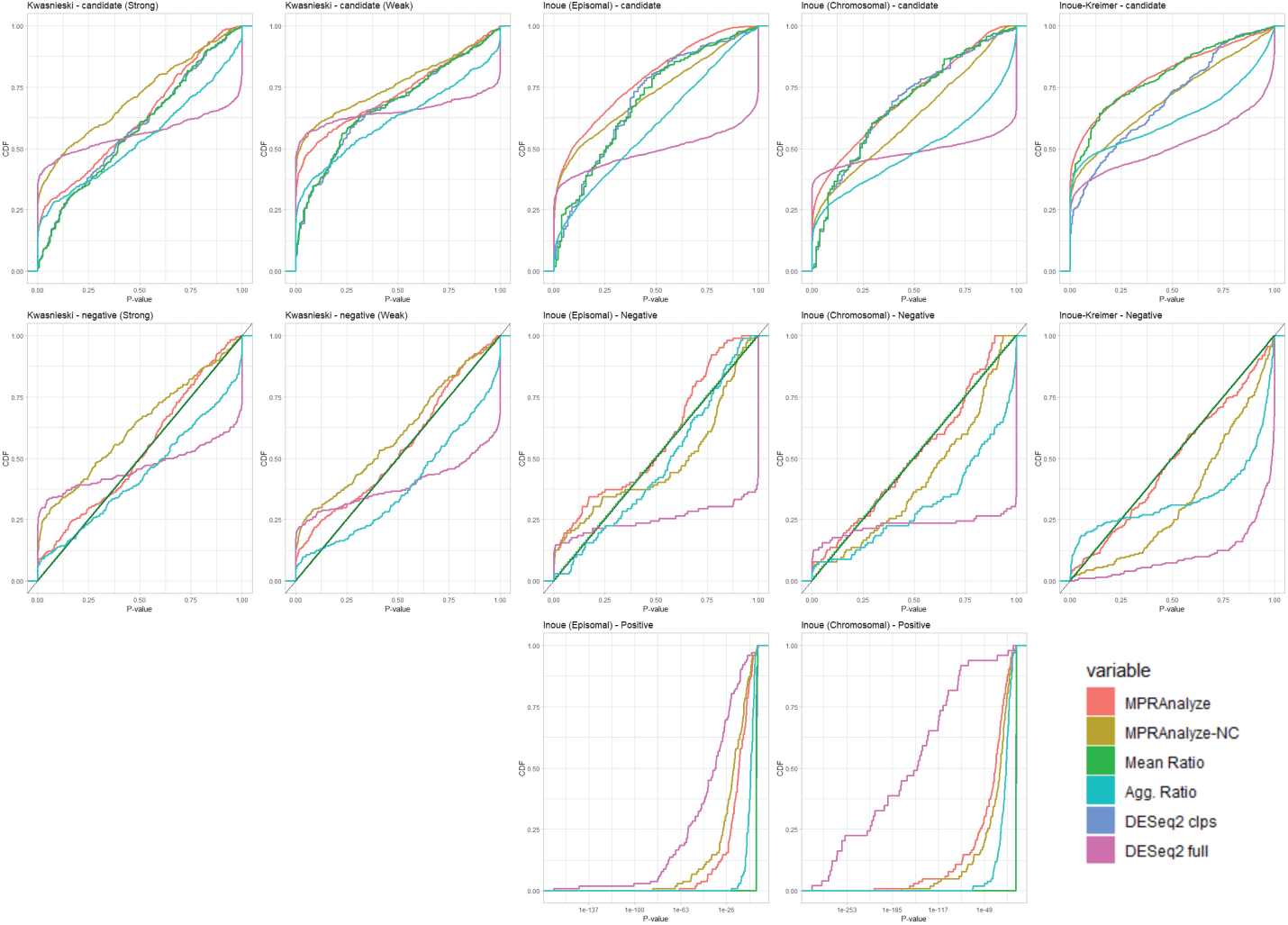
P-value CDF of classification analysis of enhancers for each dataset, stratified by enhancer type. top panels are candidate enhancers; middle are negative controls, with a reference line of the theoretical uniform CDF; bottom are positive controls, displayed in log-scale to make them more informative.

**Supplemental Figure 6:**
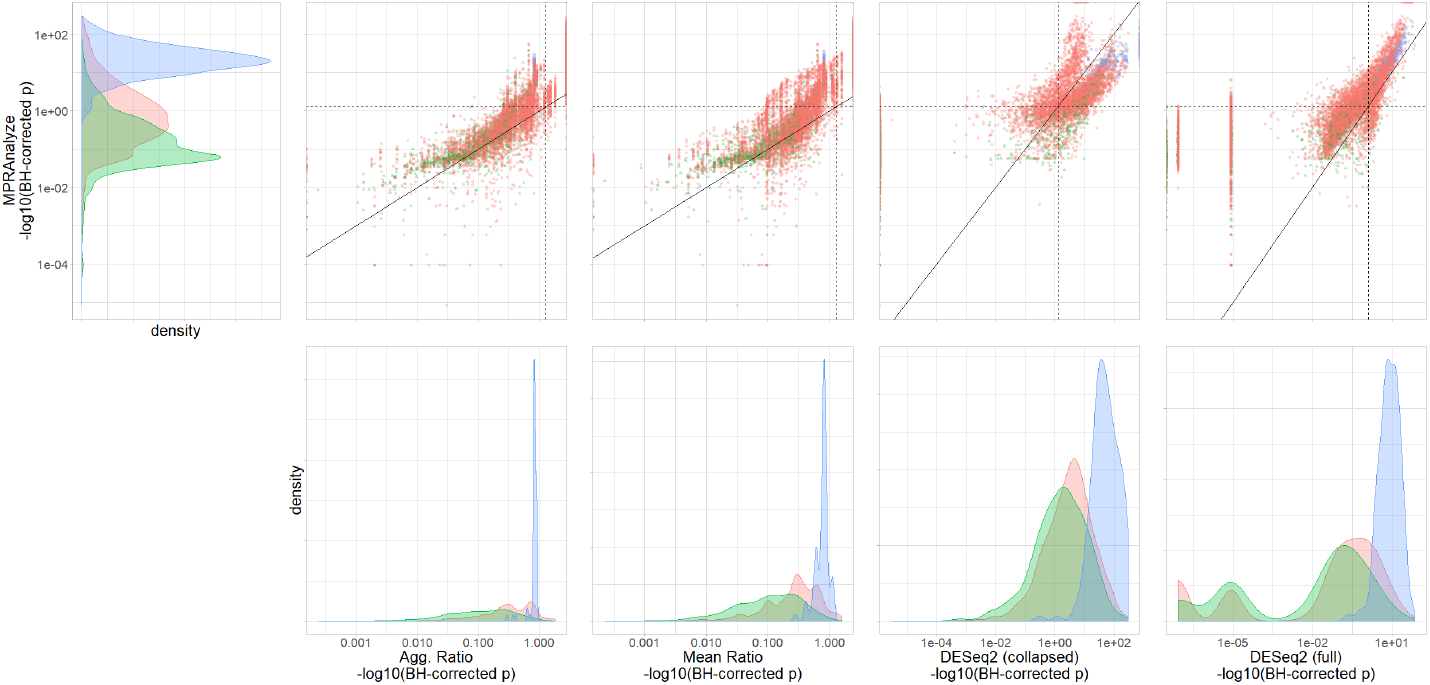
Distribution of BH-corrected P-values of all methods. Analysis was performed separately for each dataset, then P-values were combined. Positive controls are from the Inoue episomal and chromosomal datasets only (blue). Candidates (red) and negative controls (green) are from the all datasets. Continuous line is a the identity line. Dotted lines are set at 0.05

**Supplemental Figure 7:**
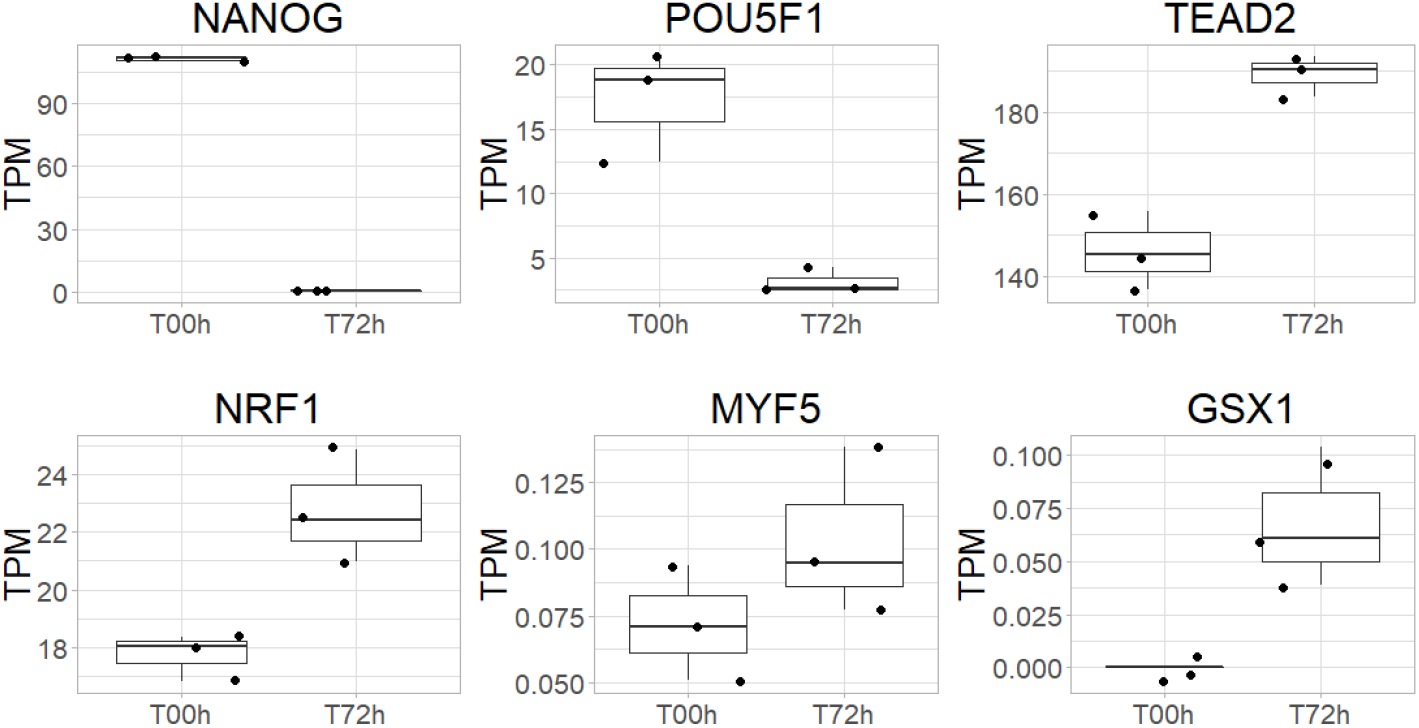
RNA expression levels for transcription factors with enriched binding sites in differentially active enhancers. NANOG and POU5F1, enriched among enhancers that reduce in activity over time, have an expected corresponding reduction in expression over time. TEAD2 and NRF1 are enriched in increasing-activity enhancers according to MPRAnalyze’s results and indeed show a corresponding increase in expression. MYF5 and GSX1 are enriched in increasing-activity enhancers according to mpralm, but not according to MPRAnalyze.

## Methods

### Dataset Collection and Processing

For all datasets included in this paper, we relied on the processing and filtering performed by the authors of the original papers. This ensures that MPRAnalyze’s performance isn’t reflecting any favorable processing steps we chose. **Kwasnieski:** The study[10] measured the activity of potential regulatory regions in K562 cells. Regions were selected according to ENCODE annotations of four groups: enhancers; weak enhancers; repressed enhancers; enhancers active in ESCs. The repressed and ESC-annotated nhancers were used as controls, and were excluded from the analysis after library size normalization factors were computed. In addition to control classes, each class had internal sets of scrambled sequences used as negative controls, which were used as controls in our analyses. **Inoue:** The study[9] compared activity in HepG2 cells of liver enhancers that were either episomal or chromosomally integrated using a lentivirus (lentiMPRA). While the study is comparative, the comparison is not between biological conditions and the results are therefore difficult to validate or interpret. We therefore decided to use the data as two separate quantification datasets. The datasets were analyzed together to better account for batch and barcode-specific effects, and *α* estimates were extracted from the joint model for each condition separately. **Inoue-Kreimer:** The study[14] identified enhancers with temporal activity over the first 72 hours after neural induction. lentiMPRA was performed in 7 timepoints (0, 3, 6, 12, 24, 48 and 72 hours after undiction). For the purpose of our analysis, we used only the data from the first timepoint in the quantification and classification analyses, and timepoints 0 and 72 hours for the comparative analysis.

### Computing Transcription Rate Estimates

All transcription rate estimates were computed for library size normalized MPRA data, using upper quartile normalization to compute size factors. **MPRAnalyze’s** *α* was computed for each dataset using the quantification analysis (See supplemental methods). Across datasets, batch and barcode-level effects were modelled in the nested DNA model, but excluded from the RNA model design. This allows MPRAnalyze to model nuisance effects but asserts that all barcodes associated with a single enhancer must share the same transcription rate. The **Mean Ratio** was computed using only pairs of observations that are both positive,so for each enhancer: *S* = {*i* ∈ [*n*] |*R_i_* ≠ 0, *D_i_* ≠ 0}, then: 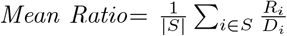.Similarly, the **Aggregated Ratio** was computed using only positive observations, without requiring that both measurements of the pair are positive, so for each enhancer: *S_R_* = {*i*|*R_i_* ≠ 0}, *S_D_* = {*i*|*D_i_* ≠ 0}, then 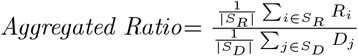.

### Subsampling analysis

For the subsampling analysis, barcodes were sampled down to varying levels (for Inoue datasets: 15, 30, 45, 60, 75, 90 out of the total 100 barcodes; for Inoue-Kreimer: 15, 30, 45, 60, 75 of the total 90 barcodes). The analysis uses three independent replicates of this down-sampling process, so overall for each enhancer we get a set of 3 × *K* estimates at various numbers of available barcodes, where *K* = 6 for the Inoue datasets and *K* = 5 for Inoue-Kreimer. The analyses were done on the entire down-sampled dataset in a single run and included the original data as well as the reduced-barcodes data, to neutralize any effect that the library size correction might have on the estimates.

### Simulating MPRA data

MPRA data was simulated by generating random coefficients for the nested GLM construction that MPRAnalyze uses. The *latent (true)* DNA and RNA counts were generated directly from the model, then log-normal noise was added to the latent counts to get the *observed* counts. Formally:

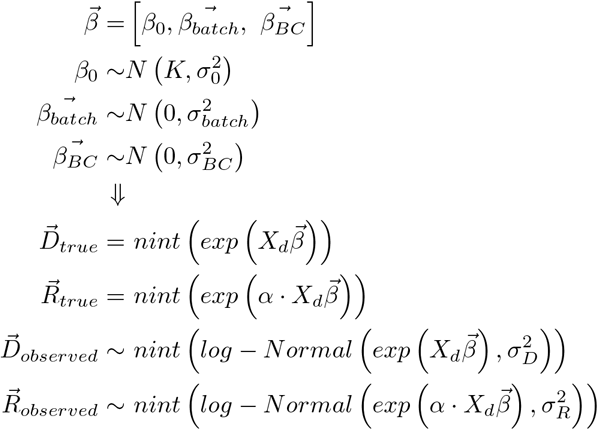

where *K* controls the intercept term for the construct distribution, the variance of which is 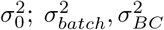 control the size of batch and barcode effects, respectively; 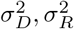 determine the noise levels added to the data; *nint* is the nearest-integer function, using base R’s *round* function. An implementation of this simulation process ins included in the MPRAnalyze package.

Noise was generated using log-normal noise instead of Gamma/Negative Binomial to avoid generating data directly from MPRAnalyze’s model, which might bias the results.

Simulated data in this manuscript was generated with 3 batches, varying numbers of barcodes, *K* = 5, and *σ*_0_ = *σ_batch_* = *σ_BC_* = *σ_D_* = *σ_R_* = 0.5.

### Transcription Factor Binding Site enrichment analysis

The transcription factor binding site enrichment analysis was performed using the binary binding matrix computed by Inoue & Kreimer et al. [14], with each entry indicating the potential for binding (motif-based binding prediction using Fimo[21], *FDR* < 10^−4^) or overlap with transcription factor ChIP-seq peaks from publicly available data[22, 23].

## References

[1] Wolfgang Huber et al. “Orchestrating high-throughput genomic analysis with Bioconductor”. en. In: Nat. Methods 12.2 (Feb. 2015), pp. 115–121. issn: 1548-7091, 1548-7105. doi: 10.1038/nmeth.3252. url: http://dx.doi.org/10.1038/nmeth.3252.

[2] Lucia A Hindorff et al. “Potential etiologic and functional implications of genome-wide association loci for human diseases and traits”. en. In: Proc. Natl. Acad. Sci. U. S. A. 106.23 (June 2009), pp. 9362–9367. issn: 0027-8424, 1091-6490. doi: 10.1073/pnas.0903103106. url: http://dx.doi.org/10.1073/pnas.0903103106.

[3] Navneet Matharu and Nadav Ahituv. “Minor Loops in Major Folds: Enhancer-Promoter Looping, Chromatin Restructuring, and Their Association with Transcriptional Regulation and Disease”. en. In: PLoS Genet. 11.12 (Dec. 2015), e1005640. issn: 1553-7390, 1553-7404. doi: 10.1371/journal.pgen.1005640. url: http://dx.doi.org/10.1371/journal.pgen.1005640.

[4] Valer Gotea et al. “Homotypic clusters of transcription factor binding sites are a key component of human promoters and enhancers”. en. In: Genome Res. 20.5 (May 2010), pp. 565–577. issn: 1088-9051, 1549-5469. doi: 10.1101/gr.104471.109. url: http://dx.doi.org/10.1101/gr.104471.109.

[5] Menno P Creyghton et al. “Histone H3K27ac separates active from poised enhancers and predicts developmental state”. en. In: Proc. Natl. Acad. Sci. U. S. A. 107.50 (Dec. 2010), pp. 21931–21936. issn: 0027-8424, 1091-6490. doi: 10.1073/pnas.1016071107. url: http://dx.doi.org/10.1073/pnas.1016071107.

[6] Nathaniel D Heintzman et al. “Histone modifications at human enhancers reflect global cell-type-specific gene expression”. en. In: Nature 459.7243 (May 2009), pp. 108–112. issn: 0028-0836, 1476-4687. doi: 10.1038/nature07829. url: http://dx.doi.org/10.1038/nature07829.

[7] Axel Visel et al. “ChIP-seq accurately predicts tissue-specific activity of enhancers”. en. In: Nature 457.7231 (Feb. 2009), pp. 854–858. issn: 0028-0836, 1476-4687. doi: 10.1038/nature07730. url: http://dx.doi.org/10.1038/nature07730.

[8] Diego Villar et al. “Enhancer evolution across 20 mammalian species”. en. In: Cell 160.3 (Jan. 2015), pp. 554–566. issn: 0092-8674, 1097-4172. doi: 10.1016/j.cell.2015.01.006. url: http://dx.doi.org/10.1016/j.cell.2015.01.006.

[9] Fumitaka Inoue et al. “A systematic comparison reveals substantial differences in chromosomal versus episomal encoding of enhancer activity”. en. In: Genome Res. 27.1 (Jan. 2017), pp. 38–52. issn: 1088-9051, 1549-5469. doi: 10.1101/gr.212092.116. url: http://dx.doi.org/10.1101/gr.212092.116.

[10] Jamie C Kwasnieski et al. “High-throughput functional testing of ENCODE segmentation predictions”. en. In: Genome Res. 24.10 (Oct. 2014), pp. 1595–1602. issn: 1088-9051, 1549-5469. doi: 10. 1101/ gr. 173518. 114. url: http://dx.doi.org/10.1101/gr.173518.114.

[11] Ryan Tewhey et al. “Direct Identification of Hundreds of Expression-Modulating Variants using a Multiplexed Reporter Assay”. en. In: Cell 165.6 (June 2016), pp. 1519–1529. issn: 0092-8674, 1097-4172. doi: 10. 1016/j.cell.2016.04.027. url: http://dx.doi.org/10.1016/j.cell.2016.04.027.

[12] Jacob C Ulirsch et al. “Systematic Functional Dissection of Common Genetic Variation Affecting Red Blood Cell Traits”. en. In: Cell 165.6 (June 2016), pp. 1530–1545. issn: 0092-8674, 1097-4172. doi: 10.1016/j.cell.2016.04.048. url: http://dx.doi.org/10.1016/j.cell.2016.04.048.

[13] Susan Q Shen et al. “Massively parallel cis-regulatory analysis in the mammalian central nervous system”. en. In: Genome Res. 26.2 (Feb. 2016), pp. 238–255. issn: 1088-9051, 1549-5469. doi: 10.1101/gr.193789.115. url: http://dx.doi.org/10.1101/gr.193789.115.

[14] Fumitaka Inoue et al. “Massively parallel characterization of regulatory dynamics during neural induction”. en. July 2018. url: https://www.biorxiv.org/content/early/2018/07/16/370452.

[15] Michael I Love, Wolfgang Huber, and Simon Anders. “Moderated estimation of fold change and dispersion for RNA-seq data with DESeq2”. en. In: Genome Biol. 15.12 (2014), p. 550. issn: 1465-6906. doi: 10.1186/ s13059-014-0550-8. url: http://dx.doi.org/10.1186/s13059-014-0550-8.

[16] Cynthia A Kalita et al. “QuASAR-MPRA: Accurate allele-specific analysis for massively parallel reporter assays”. en. In: Bioinformatics (Sept. 2017). issn: 1367-4803, 1367-4811. doi: 10.1093/bioinformatics/btx598. url: http://dx.doi.org/10.1093/bioinformatics/btx598.

[17] Leslie Myint et al. “Linear models enable powerful differential activity analysis in massively parallel reporter assays”. en. Sept. 2017. url: https://www.biorxiv.org/content/early/2017/09/30/196394.

[18] Kotaro J Kaneko et al. “Transcription factor TEAD2 is involved in neural tube closure”. en. In: Genesis 45.9 (Sept. 2007), pp. 577–587. issn: 1526-954X. doi: 10.1002/dvg.20330. url: http://dx.doi.org/10.1002/dvg.20330.

[19] Wen-Teng Chang et al. “A novel function of transcription factor alpha-Pal/NRF-1: increasing neurite outgrowth”. en. In: Biochem. Biophys. Res. Commun. 334.1 (Aug. 2005), pp. 199–206. issn: 0006-291X. doi: 10.1016/j.bbrc.2005.06.079. url: http://dx.doi.org/10.1016/j.bbrc.2005.06.079.

[20] Gil Stelzer et al. “The GeneCards Suite: From Gene Data Mining to Disease Genome Sequence Analyses”. en. In: Curr. Protoc. Bioinformatics 54 (June 2016), pp. 1.30.1–1.30.33. issn: 1934-3396, 1934-340X. doi: 10.1002/cpbi.5. url: http://dx.doi.org/10.1002/cpbi.5.

[21] Charles E Grant, Timothy L Bailey, and William Stafford Noble. “FIMO: scanning for occurrences of a given motif”. en. In: Bioinformatics 27.7 (Apr. 2011), pp. 1017–1018. issn: 1367-4803, 1367-4811. doi: 10.1093/bioinformatics/btr064. url: http://dx.doi.org/10.1093/bioinformatics/btr064.

[22] Casey A Gifford et al. “Transcriptional and epigenetic dynamics during specification of human embryonic stem cells”. en. In: Cell 153.5 (May 2013), pp. 1149–1163. issn: 0092-8674, 1097-4172. doi: 10.1016/j.cell.2013.04.037. url: http://dx.doi.org/10.1016/j.cell.2013.04.037.

[23] Alexander M Tsankov et al. “Transcription factor binding dynamics during human ES cell differentiation”. en. In: Nature 518.7539 (Feb. 2015), pp. 344–349. issn: 0028-0836, 1476-4687. doi: 10.1038/nature14233. url: http://dx.doi.org/10.1038/nature14233.

