## Supplemental Methods - model for "MPRAnalyze - A statistical framework for Massively Parallel Reporter Assays"

### Supplementary Methods: MPRAnalyze model description

### Introduction

MPRAnalyze aims to infer the true induced transcription rate $\alpha$ for each of the candidate enhancers included in a given MPRA dataset, by modeling the raw counts and avoiding ratio-based summary statistics.

Formally, the MPRA data for a given enhancer can be denoted as $\left(\vec{d}, \vec{r}\right)$ where $\vec{d} = \begin{bmatrix} d_1 \\ \vdots \\ d_m \end{bmatrix}$ and $\vec{r} = \begin{bmatrix} r_1 \\ \vdots \\ r_n \end{bmatrix}$, Where $n$ is the number of available RNA observations and $m$ is the number of available DNA observations. Note that ideally, the RNA and DNA libraries would both be from the same post-transduction populations, which would allow complete modeling of technical and confounding effects in all layers of the model. However, since many MPRA libraries sequences the DNA from the pre-transduction plasmid population, MPRAnalyze supports both options.

### Regression Model

MPRAnalyze assumes a linear relationship between RNA and DNA, an assumption easily checked and mostly supported by MPRA data. Formally, we assume that for some transcription rate $\alpha$, the true DNA and RNA measures follow a linear relationship with slope $\alpha$:

$$RNA = \alpha \cdot DNA$$

However, the data is usually generated by a set of different distributions which depend on wanted sources of variation like different cell types or cellular environments, or unwanted sources of variation like batch effects or barcode-specific effects. To enable controlling these factors in the analysis, MPRAnalyze assumes all effects are independent and additive in log space and implements a flexible and extendable regression model.

We construct a separate regression model for the DNA and RNA models, and fit them by likelihood maximization.

Formally, we model each observation as follows:

$$\hat{d}_i = \left(\prod_{\ell} \exp\left(\beta_{\ell(i)}\right)\right) \cdot S_{d,i} \Rightarrow \log\left(\hat{d}_i\right) = \sum_{\ell} \beta_{\ell(i)} + \log\left(S_{d,i}\right)$$

$$\hat{r}_i = \hat{\alpha}\hat{d}_i \cdot S_{r,i} = \left(\prod_{m} \exp\left(\gamma_{m(i)}\right)\right) \cdot \left(\prod_{\ell} \exp\left(\beta_{\ell(i)}\right)\right) \cdot S_{r,i} \Rightarrow$$

$$\Rightarrow \log\left(\hat{r_i}\right) = \sum_{\ell} \beta_{\ell(i)} + \sum_{m} \gamma_{m(i)} + \log\left(S_{r,i}\right)$$

Where $S_{d,i}, S_{r,i}$ are library depth normalization factors, $\ell, m$ relate the observation index to the appropriate batch factor, and $\beta, \gamma$ are the set of parameters to optimize.

In this framework, $\prod_{\ell} \beta_{\ell(i)}$ is the estimate of the true number of DNA molecules, and $\prod_{m} \gamma_{m(i)}$ is the estimate of $\alpha$, the transcription rate. Note that library depth factors are purely technical and therefore $S_{d,i}$ is only corrected for when fitting the DNA model and not included in the RNA model, and that these factors can be estimated externally. Fitting the parameters across the observations can be done by matrix multiplication with binary design matrices $X_d,\ X_r$, simplifying to:

$$\log\left(\vec{\hat{d}}\right) = X_d\vec{\beta} + \log\left(\vec{S_d}\right)$$

$$\log\left(\vec{\hat{r}}\right) = X_d\vec{\beta} + X_r\vec{\gamma} + \log\left(\vec{S_r}\right)$$

We then optimize these parameters by optimizing the likelihood function of the data, conditioned on some distributional assumptions (detailed below). Formally we use the vector of estimates $\vec{\hat{d}}, \vec{\hat{r}}$ as estimators for the first moment of the distribution the observations were generated by. The exact likelihood computation depends on the distributional model, but conceptually we get:

$$\begin{aligned}
\left(\beta^{MLE}, \gamma^{MLE}\right) &= \arg\max_{\vec{\beta},\vec{\gamma}} \left\{\mathcal{L}_D\left(\vec{d} \mid \vec{\hat{d}}\right) \cdot \mathcal{L}_R\left(\vec{r} \mid \vec{\hat{r}}\right)\right\} \\
&= \arg\max_{\vec{\beta},\vec{\gamma}} \left\{\mathcal{L}_D\left(\vec{d} \mid \vec{\hat{d}}\right) \cdot \mathcal{L}_R\left(\vec{r} \mid \vec{\hat{d}} \cdot \hat{\alpha} \cdot \vec{S_r}\right)\right\} \\
&= \arg\max_{\vec{\beta},\vec{\gamma}} \left\{\ell_D\left(\vec{d} \mid\ \exp\left(X_d\vec{\beta} + \log\left(\vec{S_d}\right)\right)\right) + \ell_R\left(\vec{r} \mid\ \exp\left(X_d\vec{\beta} + X_r\vec{\gamma} + \log\left(\vec{S_r}\right)\right)\right)\right\} \\
&= \arg\max_{\vec{\beta},\vec{\gamma}} \left\{\ell_D\left(\vec{d} \mid\ \exp\left(X_d\vec{\beta}\right) \cdot \vec{S_d}\right) + \ell_R\left(\vec{r} \mid\ \exp\left(X_d\vec{\beta}\right) \cdot \exp\left(X_r\vec{\gamma}\right) \cdot \vec{S_r}\right)\right\}
\end{aligned}$$

With $\mathcal{L}_D, \mathcal{L}_R$ are the likelihood functions of the DNA, RNA distributions (respectively), and $\ell_D, \ell_R$ their respective log-likelihood functions.

### Distributional Models

MPRAnalyze models the generation of the data as:

$$\vec{r} = \alpha\vec{d}$$

The main complexity of the model therefore stems from the RNA observations being a convolution of the scaled DNA distribution and the RNA measurement distribution. We offer two ways to handle this:

### 1 Closed Form Solution

Some distributions provide a closed form for this scenario. MPRAnalyze uses this by assuming a Gamma-Poisson model: we assume the DNA model follows a Gamma distribution: $\vec{d} \sim Gamma\left(k, b\right)$ and that $\vec{r}|\vec{d} \sim Poisson\left(\alpha \cdot \vec{d}\right)$. The resulting likelihood for the RNA model has a closed form as a Negative-Binomial distribution, which is a well established approximation of sequencing data. Formally:

$$\begin{aligned}
\vec{d} &\sim Gamma\left(k,\ \beta\right) \\
\Rightarrow \alpha \cdot \vec{d} &\sim Gamma\left(k,\ \frac{\beta}{\alpha}\right) \\
\vec{r}|\vec{d} &\sim Poisson\left(\alpha \cdot \vec{d}\right) \\
\Rightarrow \vec{r} &\sim Negative-Binomial\left(\mu = \frac{\alpha \cdot k}{\beta}, \psi = k\right)
\end{aligned}$$

To incorporate the regression scheme described above in this setting, we share the dispersion parameter $a$ between the DNA and RNA models. The DNA regression model is the estimate for $\left(\frac{1}{b}\right)$, and the RNA regression model is the estimate for $\alpha$. Therefore, the MLE estimates are:

$$\begin{aligned}
\left(\beta^{MLE}, \gamma^{MLE}, a^{MLE}\right) &= \arg\max_{\vec{\beta}, \vec{\gamma}, \hat{\alpha}} \left\{ \mathcal{L}_{Gamma}\left(\vec{d} \mid a = \hat{a}, b = \hat{b}\right) \cdot \mathcal{L}_{NB}\left(\vec{r} \mid \mu = \frac{\hat{a} \cdot \hat{\alpha}}{\hat{b}}, \psi = \hat{a}\right)\right\} \\
&= \arg\max_{\vec{\beta}, \vec{\gamma}, \hat{\alpha}} \left\{ \ell_{Gamma}\left(\vec{d} \mid a = \hat{a}, b = \left(\exp\left(X_d \vec{\beta}\right) \cdot \vec{S}_D\right)^{-1}\right) \right. \\
&\left. + \ell_{NB}\left(\vec{r} \mid \mu = \hat{a} \cdot \exp\left(X_d \vec{\beta}\right) \cdot \exp\left(X_r \vec{\gamma}\right) \cdot \vec{S}_R, \psi = \hat{a}\right)\right\}
\end{aligned}$$

### 2 Point estimator

Alternatively, the model can be simplified by fitting the DNA model and collapsing it to a point estimator. Unlike the closed form solution, this means that not all of the information of the DNA model will be shared with the RNA model. However, this means that this option allows for more flexibility in the distributional assumptions, essentially allowing any pair of distributions to be used for the DNA and RNA models. This flexibility is mostly important for the restrictions imposed on the variance estimation. MPRAnalyze implements this approach in two ways, allowing for two sets of constraints on the variance.

#### 2.1 log-Normal DNA, Negative-Binomial RNA

assuming:

$$\begin{aligned}
\vec{d} &\sim log-Normal\left(m, \sigma^2\right) \\
\vec{r} &\sim Negative-Binomial\left(\mu = \alpha m,\ \psi\right)
\end{aligned}$$

Which, along with the regression scheme, translates to:

$$\left(\beta^{MLE}, \gamma^{MLE}, \sigma^{MLE}, \psi^{MLE}\right) = \arg\max_{\vec{\beta},\vec{\gamma},\hat{\sigma},\hat{\psi}} \left\{\mathcal{L}_{logNormal}\left(\vec{d} \mid m = \hat{m}, \sigma^2 = \hat{\sigma}^2\right) \cdot \mathcal{L}_{NB}\left(\vec{r} \mid \mu = \hat{m} \cdot \hat{\alpha}, \psi = \hat{\psi}\right)\right\}$$
$$= \arg\max_{\vec{\beta},\vec{\gamma},\hat{\sigma},\hat{\psi}} \left\{\ell_{logNormal}\left(\vec{d} \mid m = \exp\left(X_d\vec{\beta}\right) \cdot \vec{S}_D, \sigma^2 = \hat{\sigma}^2\right)\right.$$
$$\left. + \ell_{NB}\left(\vec{r} \mid \mu = \exp\left(X_d\vec{\beta}\right) \cdot \exp\left(X_r\vec{\gamma}\right) \cdot \vec{S_R}, \psi = \hat{\psi}\right)\right\}$$

##### 2.2 log-Normal DNA, log-Normal RNA

assuming:

$$\vec{d} \sim log - Normal\left(m, \sigma_d^2\right)$$
$$\vec{r} \sim log - Normal\left(m \cdot \alpha, \sigma_r^2\right)$$

which, along with the regression scheme, translates to:

$$\left(\beta^{MLE}, \gamma^{MLE}, \sigma_d^{MLE}, \sigma_r^{MLE}\right) = \arg\max_{\vec{\beta},\vec{\gamma},\hat{\sigma_d},\hat{\sigma_r}} \left\{\mathcal{L}_{logNormal}\left(\vec{d} \mid \mu = \hat{m}, \sigma^2 = \hat{\sigma_d}^2\right) \cdot \mathcal{L}_{logNormal}\left(\vec{r} \mid \mu = \hat{m} \cdot \hat{\alpha}, \sigma^2 = \hat{\sigma_r}^2\right)\right\}$$
$$= \arg\max_{\vec{\beta},\vec{\gamma},\hat{\sigma_d},\hat{\sigma_r}} \left\{\ell_{logNormal}\left(\vec{d} \mid \mu = \exp\left(X_d\vec{\beta}\right) \cdot \vec{S}_D, \sigma^2 = \hat{\sigma_d}^2\right)\right.$$
$$\left. + \ell_{logNormal}\left(\vec{r} \mid \mu = \exp\left(X_d\vec{\beta}\right) \cdot \exp\left(X_r\vec{\gamma}\right) \cdot \vec{S_R}, \sigma^2 = \hat{\sigma_r}^2\right)\right\}$$

### Hypothesis testings

MPRAnalyze implements a variety of hypothesis testing options to accommodate the variety of different MPRA experimental design settings. The two major types of analyses MPRA facilitates are quantitative analysis: determining the level of transcriptional activity induced by each candidate enhancer; and comparative analysis: identifying differential activity of a candidate enhancer between biological conditions. MPRAnalyze provides a robust statistical framework that enables addressing these questions while leveraging the existence of negative controls (scramble sequences) in the experiment. The various use cases are summarized by the following diagram, and fully explained below:

#### Comparative Analysis

A comparative analysis is one where the leading question is about the difference in activity of a given enhancer between conditions. Examples of these include comparing enhancer activity between cell types to detect tissue-specific activity, comparing different time-points in a differentiation process and comparing activity in unstimulated cells with cells exposed to various stimuli. MPRAnalyze offers several possible settings and testing frameworks to address these kinds of questions.

###### Coefficient-based Testing

A coefficient-based testing framework addresses these comparisons by directly testing the regression coefficients for significane, using a Wald test. This approach enables testing multiple different hypotheses using the same model fit, which can be useful in some cases. For example, if effects of different stimuli are to be tested, a single model can be fit to the entire dataset, then testing the effect of each stimulus is simply computing the Wald-test for the appropriate regression coefficient, with the unstimulated samples as the reference.

##### Likelihood-Ratio Testing

A likelihood Ratio test lacks the post-fitting flexibility of a coefficient-based testing, but allows more complex questions to be asked. For instace, looking at a time-series dataset of a differentiation process, one might ask "does this enhancer change their activity over time". This question can be difficult to address with coefficient-based testing, since each single coefficient only corresponds to one time point. However, this can easily be addressed with a likelihood-ratio test. This is done by fitting two models: a 'full' model with the complete design including the conditional factor (in this case, the factors that identify the time-point each data-point is associated with), and another model - a 'reduced' model - that is identical to the full model in everything expect that it lacks this factor of interest. Since the reduced space is a sub-space of the full space (in which the time-related coefficients are 0), these models are nested and a standard Likelihood-Ratio Test can be used to test for statistical significance.

### Classification Analysis

A classification analysis address the question of the overall degree of activity of a given enhancer. Note that addressing this question without an established baseline for the basal transcription rate is difficult, and the power and interpretability of these analyses are very limited in the absence of negative controls. MPRAnalyze performs classification by testing each enhancer's estimated transcription rate, $\alpha$, against the null distribution. This null distribution is based on the *alpha*s of the negative controls if available, and on the bottom half of the total distribution otherwise, under the implicit assumption that active enhancers will be more active than the median enhancer. In either case, MPRAnalyze computes both a Z-score and a MAD-score, which is a median-based variant of the Z-score. These scores are normally distributed under the null, and we can therefore compute p-values from the standard normal distribution.

#### Classification with Controls

Negative controls can be used directly to establish the null distribution, then computing the scores within the null. Formally, for $\vec{\alpha_c} = (\alpha_{c_1}, \ldots, \alpha_{c_\ell})$ the estimated rates for the $\ell$ controls included in the experiment:

$$\text{MAD-score}_c(\alpha_i) = \frac{\alpha_i - \text{median}(\vec{\alpha_c})}{\text{K} \cdot \text{median}(|\vec{\alpha_{c_i}} - \text{median}(\vec{\alpha_c})|)}$$

$$\text{Z-score}_c(\alpha_i) = \frac{\alpha_i - \text{mean}(\vec{\alpha_c})}{\text{std}(\vec{\alpha_c})}$$

Additionally, MPRAnalyze also calculates the empirical p-value:

$$P(\alpha_i) = \frac{1}{\ell} \sum_{j=1}^{\ell} 1\left(\alpha_i > \alpha_{c_j}\right)$$

##### Classification without Controls

Without the advantage of negative controls to establish the null transcription rate, MPRAnalyze empirically determines the distributional properties of the null set. For this, MPRAnalyze relies on three assumptions: (1) the data contains both active and inactive enhancers; (2) active enhancers generally have a higher estimated rate of transcription than inactive enhancers; (3) not all active enhancers are equally active, but all inactive enhancers are equally inactive, meaning generated by the same distribution determined by the properties of the promoter used. MPRAnalyze empirically estimates the mode of the distribution, by taking the center of the window which contains the maximal number of samples, with a window size set to be 0.02 of the total range of values in the sample. This mode is assumed to be the center of the null distribution, due to the third assumption. Then samples that are lower from the mode are then defined as the null set, and are used to estimate the width of the distribution, assuming the null is symmetric with respect to the mode.

The calculation of the scores is done using the null set:

$$\text{MAD-score}_{\text{null}} = \frac{\alpha_i - \text{mode}\left(\vec{\alpha}\right)}{\text{K} \cdot \text{median}\left(\text{mode}\left(\vec{\alpha}\right) - \vec{\alpha_{\text{null}}}\right)}$$

$$\text{Z-score}_{\text{null}} = \frac{\alpha_i - \text{mode}\left(\vec{\alpha}\right)}{\sqrt{\text{mean}\left(\left(\text{mode}\left(\vec{\alpha}\right) - \vec{\alpha_{\text{null}}}\right)^2\right)}}$$

### Incorporating Negative Controls

MPRAnalyze enables incorporating negative controls into the model to correct systemic bias in the data. Such bias often stems from inherent biological differences between condition that are not related to the regulatory mechanism studied, e.g cellular size differences. Including negative controls in the experiment, preferably by including randomized "scrambled" sequences in the experiment, allows quantifying and correcting these effects. This can be done in several ways, all of which involve some manner of fitting a joint model for control enhancers.

All negative controls are assumed to be generated by the same, baseline, RNA model. However, since there's no reason to assume this has any effect on transfection efficiency, a separate DNA model is still fit to each control enhancer. Formally, for $n$ control enhancers, the joint model is:

$$\left(\beta_1^{MLE}, \ldots, \beta_n^{MLE}, \gamma^{MLE}\right) =$$
$$\arg \max_{\beta_1, \ldots, \beta_n, \gamma} \left\{ \sum_{i=1}^{n} \ell_D \left( \vec{d} \mid \exp\left(X_d \vec{\beta_i}\right) \cdot \vec{S_d},\ \psi_D \right) + \sum_{i=1}^{n} \ell_R \left( \vec{r} \mid \exp\left(X_d \vec{\beta_i}\right) \cdot \exp\left(X_r \vec{\gamma}\right) \cdot \vec{S_r},,\ \psi_R \right) \right\}$$

Fitting a joint models of or with controls can be used in several ways to account for systemic bias in the data.

###### Control-based Correction Factors

The joint control model described above can be used to generate normalization factors based on systemic bias reflected in the

A separate model can be fit for the controls, and correction factors extracted from it and included in the general model. Similar to library-depth correction, this would amount to an additional scaling factor added to the regression scheme that corrects the inherent bias in the system as measured by the negative controls.

### Theoretical Supplements

#### Gamma-Poisson convolution proof

Proof of the closed form relationship of the likelihood of the RNA model in the Gamma-Poisson model:

Let $D \sim Gamma\left(k, \beta\right)$ and $R|D \sim Poisson\left(\lambda = \alpha D\right)$.

First, we'll note that $\alpha D \sim Gamma\left(k, \frac{\beta}{\alpha}\right)$, and therefore:

$$
\begin{aligned}
P\left(R = x\right) &= \int_0^\infty P\left(R = x|\lambda\right) P_{\alpha D}\left(\lambda\right) d\lambda \\
&= \int_0^\infty \frac{\lambda^x e^{-\lambda}}{x!} \cdot \frac{\left(\frac{\beta}{\alpha}\right)^k}{\Gamma\left(k\right)} \cdot \lambda^{k-1} e^{-\frac{\beta\lambda}{\alpha}} d\lambda \\
&= \frac{\left(\frac{\beta}{\alpha}\right)^k}{\Gamma\left(k\right) \cdot x!} \int_0^\infty \lambda^{k+x-1} e^{-\lambda\left(\frac{\beta}{\alpha}+1\right)} d\lambda \\
&= \frac{\left(\frac{\beta}{\alpha}\right)^k}{\Gamma\left(k\right) \cdot x!} \cdot \left(1 + \frac{\beta}{\alpha}\right)^{-(k+x)} \Gamma\left(k + x\right) \\
&= \frac{\Gamma\left(k + x\right)}{\Gamma\left(k\right) \cdot x!} \cdot \left(\frac{\frac{\beta}{\alpha}}{1 + \frac{\beta}{\alpha}}\right)^k \cdot \left(\frac{1}{1 + \frac{\beta}{\alpha}}\right)^x \\
\Rightarrow & R \sim NB\left(\mu = \frac{\alpha \cdot k}{\beta},\ \psi = k\right)
\end{aligned}
$$
